# Chemovaccination with a novel antimalarial targeting the late liver stage induces durable immunity against malaria

**DOI:** 10.1101/2025.07.22.666032

**Authors:** Ryan W.J. Steel, Yu Cheng Chua, Sabrina Caiazzo, Eva Hesping, Daniel Fernandez-Ruiz, Lauren E. Holz, William R. Heath, John A. McCauley, David B. Olsen, Justin A. Boddey

## Abstract

Vaccination with *Plasmodium falciparum*, the most lethal malaria parasite, using sporozoites that arrest during liver stage infection either by irradiation, genetic attenuation or chemotherapy have been developed, with late liver stage arrest providing very high efficacy. Such vaccines require complex manufacture, deployment and intravenous administration. Here, we report an alternative strategy of chemo-attenuation of malaria parasites at the late liver stage using first-in-class antimalarials under clinical development that target the parasite aspartyl proteases plasmepsin IX and X. A single low-dose infection with virulent *Plasmodium berghei* sporozoites followed by drug treatment cleared infection by producing <u>c</u>hemo-<u>a</u>ttenuated liver <u>m</u>erozoites (CALM) that induced sterile immunity in mice for up to 21 months. Protection arose from humoral responses to circumsporozoite protein and robust CD8^+^ T cell responses, including liver-resident memory cells reactive to diverse antigens including SERA1 and RPL6. Drug treatment also attenuated the human pathogen *P. falciparum* by preventing liver merozoites from infecting human erythrocytes in humanized chimeric liver mice, confirming that the mechanism of liver-stage merozoite attenuation (ie, CALM) via inhibition of plasmepsins IX and X is conserved, likely due to conservation of binding site amino acids of both proteases across the *Plasmodium* genus. Therefore, plasmepsin IX/X-targeting antimalarials offer a new approach to achieving late liver stage arrest against all circulating *Plasmodium* species and strains. This study establishes the basis for clinical trials assessing CALM for chemoprevention and chemovaccination against diverse *Plasmodium* species to advance new therapeutic strategies in malaria control. It also suggests the prospect of chemovaccination by natural exposure to mosquito-borne parasites if development of a long-acting injectable formulation of plasmepsin IX/X inhibitors proves feasible.

## Main text

Malaria continues to be a major cause of morbidity and mortality globally. In 2023, 263 million cases and 597,000 deaths were reported worldwide, an increase of 11 million cases from 2022 (*1*). Since 2000, approximately 2.2 billion cases of malaria have been averted due to implementation of control measures but control efforts are faltering, especially in Africa, where in 2023, 94% of cases and 95% of deaths from malaria occurred, and 76% of deaths occurred in children aged younger than 5 years old (*1*). Eliminating malaria requires multi-faceted approaches that combine interventions to address the complex nature of this disease.

Sporozoites are the lifecycle stage of malaria parasites that are transmitted by the bites of infected mosquitoes. After penetrating the skin they reach the liver where they invade hepatocytes and develop for several days before undergoing schizogony resulting in tens of thousands of mature liver merozoites per infected hepatocyte (*2*). Liver merozoites emerge from hepatocytes in small vesicular packets of 100 to 200 merozoites encased within hepatocyte membrane called merosomes (*3, 4*). After exiting the liver sinusoids, merosomes have been observed in mice to migrate intact through the heart and accumulate in the lungs before disintegrating, releasing merozoites into the circulation, thereby initiating the blood stage of malaria infection (*4*).

In 2021, the first circumsporozoite protein (CSP) subunit vaccine RTS,S (Mosquirix) was recommended by the World Health Organization (WHO) (*5, 6*). Four doses of RTS,S given to children aged 5 to 17 months old had 36% efficacy against malaria over a 4 year range (*5*). More recently, a second CSP subunit vaccine with a higher density of CSP antigen, R21/Matrix-M, given three times to children 5 to 36 months of age had 75% efficacy against malaria, with a booster at 12 months required to retain efficacy (*7*) and R21/Matrix-M was recommended by the WHO in 2023 (*7, 8*). When administered just before the malaria season as 3 doses, both RTS,S and R21/Matrix-M had similar efficacy (*9*). The efficacy of RTS,S and R21 varies geographically, with higher protection observed in regions with seasonal transmission and lower parasite exposure, highlighting the need for context-specific vaccination strategies. CSP polymorphisms, natural waning of protective immunity, and low CD8^+^ T-cell responses are important issues that have arisen from clinical trials with RTS,S and R21. The development of a highly efficacious vaccine that provides robust, long-lasting and heterologous protection against diverse strains and species remains an important goal to reduce and ultimately eliminate malaria (*10*).

An alternative vaccine approach that is highly efficacious in mice and humans relies on immunization with live attenuated sporozoites. In this approach, live sporozoites given by intravenous injection or mosquito bites infect the liver and replicate within hepatocytes until infection is arrested, preventing disease while presenting thousands of potential antigens for immune education. Methods that prevent the parasites from entering the blood stage include genetic attenuation through irradiation (*11*), targeted gene disruptions (*12–15*), or the use of antimalarial chemoprophylaxis to arrest infection (*16–21*). Sporozoite vaccines that abort late during hepatocyte infection (*22–27*) are more efficacious, while also requiring lower sporozoite immunization doses than sporozoite vaccines that arrest early, such as gamma-irradiated (*11*) or early genetically attenuated (*12–15*) sporozoites. It is hypothesised that the growth of individual sporozoites into tens of thousands of liver merozoites that express a broad repertoire of liver- and blood-stage antigens result in greater immune protection (*24*). Another advantage of whole sporozoite vaccines is their ability to elicit both humoral and memory T cell responses, contributing to long-lasting protection. The production of CD8^+^ T cells including liver tissue resident memory (Trm) T cells results in optimal protection in mice and non-human primates. These cells have also been associated with protection in humans, though have been challenging to isolate in humans (*28–38*). Following sporozoite vaccination of nonhuman primates, *P. falciparum*-specific CD8^+^ T cells were approximately 100-fold higher in the liver than in the blood, implicating them in long-term sterile protection despite waning antibody levels (*33*).

A particularly efficacious vaccination strategy involves sporozoite infection under the cover of chloroquine (CQ) chemoprophylaxis (CPS), so-called chemovaccination that achieved 100% efficacy after 3 doses (*36, 39–41*). CQ allows complete sporozoite development through the liver stage before killing parasites at the subsequent blood stage after a transient period of malarial symptoms (*36, 42*). Resistance to CQ spread globally in the 1950s to 1960s, precluding its widespread use in current malaria programs, including for chemovaccination. Alternative drugs that attenuate parasites while maintaining the broad antigen repertoire of late liver stage parasites represent an attractive approach to induce this highly efficacious immunity. Although numerous alternative antimalarials have been assessed, including pyrimethamine (*41*) and azithromycin (*43*), which have liver stage activity, none have yet provided the same high efficacy of protection in clinical trials as with CQ (*21, 44*).

Recently, a genetically attenuated *P. falciparum* sporozoite vaccine that arrests at late liver-stage schizonts (named genetically attenuated 2 [GA2]), given as three immunizing doses of 50 mosquito bites 28 days apart to non-immune volunteers achieved 89% efficacy against homologous challenge 3 weeks later (*25*). GA2 achieved similar efficacy with a single immunization of 50 mosquito bites when a homologous challenge was given 6 weeks later (*26*). These trials demonstrate the significance and potency of the late liver-stage for vaccinating humans (*26*), laying the groundwork for future research in this area. A drug similarly targeting the late liver stage, mimicking the late arresting genetically modified parasite approach, represents an attractive approach to sporozoite vaccination against all circulating *Plasmodium* strains and species (*45*).

Despite their high protective efficacy, sporozoite-based malaria vaccines face hurdles in scale-up and commercial manufacturing. The need for aseptic production of live, attenuated *Plasmodium* sporozoites under stringent conditions, often involving dissection of infected mosquitoes or complex bioreactor systems, poses logistical, regulatory, and cost-related challenges that impede broad deployment in endemic settings.

Here, we identify a first-in-class picomolar potent antimalarial that produces <u>c</u>hemo-<u>a</u>ttenuated liver <u>m</u>erozoites (CALM) that cannot infect erythrocytes or cause disease. CALM are potently immunogenic, conferring durable sterile protection against lethal *P. berghei* sporozoite infections in three inbred and outbred mouse species. CALM present diverse liver- and blood-stage antigens for immune education, resulting in humoral and cellular responses that protect against future infection, both by preventing sporozoite infection of hepatocytes and by CD8^+^ T cell-dependent clearance of infected hepatocytes. CALM involves the treatment and cure of liver-stage infection with a drug class under clinical development that targets the essential proteases plasmepsin IX and X (PMIX/X), whose dual inhibition renders liver-stage merozoites, including released merosomes, noninfectious for erythrocytes (*46*). WM382 also prevented the liver-to-blood transition by *P. falciparum*, the most fatal malaria parasite, in humanized mice, thus establishing the preclinical foundation for CALM vaccination trials in humans with *P. falciparum* sporozoites. PMIX/X dual inhibitors taken orally could provide CALM vaccination using cryopreserved sporozoites as a pan-*Plasmodium* injectable vaccine, since the catalytic clefts of PMIX/X are highly conserved in all human-infective species. Moreover, CALM suggests the prospect of chemovaccination by natural exposure to vector-borne parasites if development of a long-acting injectable formulation of PMIX/X inhibitors proves feasible.

## Results

### Treatment of liver infection with WM382 vaccinates and protects mice against reinfection

In previous studies, we demonstrated that the PMIX/X dual inhibitor WM382 permitted *P. berghei* sporozoite development within hepatocytes into mature liver merozoites, with normal schizogony, cytomere formation and MSP1-positive liver stage development (*46, 47*). However, while egress of merosomes proceeded with only modest impairment, their infectivity of erythrocytes was completely inhibited by WM382, providing chemoprophylaxis against malaria (*46, 47*). Analysis of RNA sequencing and proteomic studies of *P. berghei* liver stages (*48, 49*) showed high expression of PMIX, PMX, and numerous substrates of these proteases (*46, 50–52*) within mature liver merozoites and merosomes (**Fig. 1A, B**). This included the PMX substrate subtilisin 1 (SUB1) that facilitates hepatocyte egress by *P. berghei* liver merozoites (*53*). The pre-erythrocytic expression profiles of PMIX, PMX, and various substrates provide an obvious rationale for the mechanism of WM382 activity against liver merozoites while permitting development of exoerythrocytic forms within hepatocytes until the very last step of the liver stage (*46*).

**Fig. 1.**
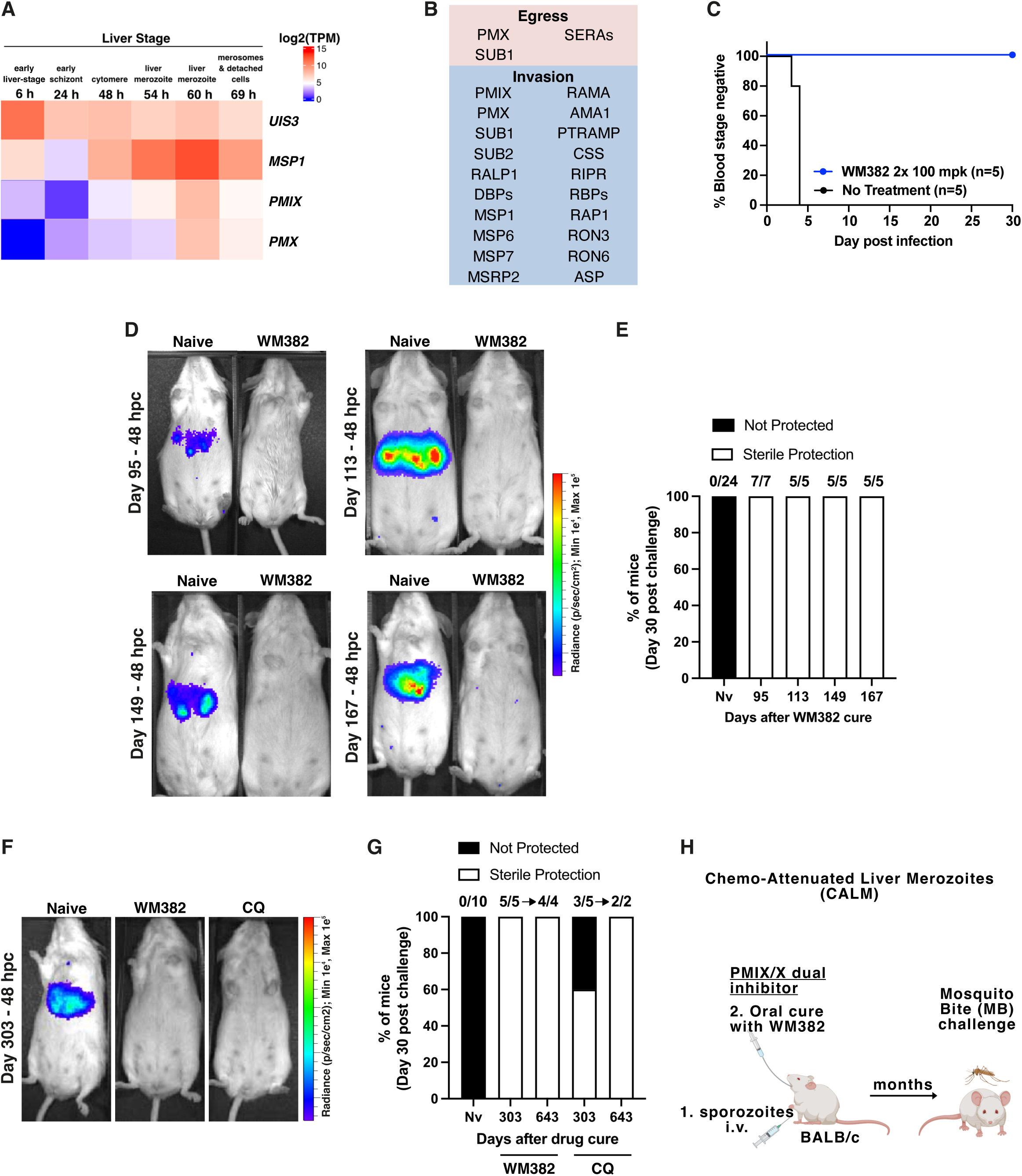
BALB/c mice with *Plasmodium* liver stage infection treated with WM382 have sterile immunity. (**A)** RNAseq heatmap of *P. berghei* liver-stage gene expression for *UIS3* (upregulated in infectious sporozoites 3; early), *MSP1* (merozoite surface protein 1; late), *PMIX* (plasmepsin IX) and *PMX (plasmepsin X).* Data mined from reference (*49*); h, hours postinfection; TPM, transcripts per million. **(B)** *P. berghei* proteins expressed in merosomes: PMIX, PMX, and various proteins and families that are known substrates (data mined from reference (*48*)). Proteins are categorised by their roles in egress and invasion of erythrocytes, while SUB1 and SERA4 are implicated in hepatocyte egress. (**C)** Sporozoite infection of BALB/c mice and treatment with 100 mg/kg (mpk) WM382 at 36- and 48-hours postinfection (hpi) prevents blood stage infection. **(D)** Sporozoite infection and WM382 treatment (see C) vaccinates and protects mice against bites from 10 mosquitoes given 95 to 167 days later, measured 48 hours postchallenge (hpc) by IVIS. **(E)** Naive mice in (D) developed lethal blood stage infections, but no mice previously treated with WM382 in (D) developed parasitemia for 30 days postchallenge, demonstrating sterile protection. **(F)** Sporozoite infection and treatment with WM382 or chloroquine (CQ) greatly reduces liver infection after mosquito bite challenge given 303 days later, measured 48 hours postchallenge (hpc) by IVIS. **(G)** Protection of mice in (F) from blood stage infection for 30 days. All mice previously treated with WM382 (5/5) and three of five mice previously treated with CQ were protected for 30 days. Surviving mice (four from WM382 group, two from CQ group) were re-challenged by mosquito bites 643 days following vaccination and remained protected for 30 days. **(H)** Intravenous (IV) infection with *P. berghei* sporozoites and treatment with WM382 at 36- and 48-hpi generates chemo-attenuated liver merozoites (CALM) that vaccinate BALB/c mice, affording sterile protection.

Prompted by prior implementation of chemoprophylaxis to attenuate sporozoite vaccines (reviewed in (*21*)), our studies of PMIX and PMX liver-stage inhibition were extended to test whether WM382 treatment resulted in an immunogenic response in animals. To this end, BALB/c mice were given a single infection of 40,000 *P. berghei* (Pb) sporozoites expressing mCherry and Luciferase (PbmChLuc) and WM382 chemoprophylaxis to treat their infection at the very late liver stage, as described previously (*46*) (**Fig. 1C**). These mice were then challenged with bites of 10 PbmChLuc infected mosquitoes 3 to 10 months (95 to 303 days) later without any further drug treatment. Since the half-life of WM382 in mice is approximately 2.5 hours, no significant drug remained in the animals at the time of challenge. All animals treated with WM382 before challenge had negligible liver infections and no parasitemia or malaria for the next 30 days indicating they were fully protected against reinfection (**Fig. 1D-G, S1A, B; Table 1 Groups 1-5**). In contrast, treatment-naive mice all developed liver infections from the mosquito bites and virulent malaria requiring euthanasia, as expected. Mice protected from challenge at 10 months (303 days) were rechallenged with mosquito bites at 21 months (643 days) without further drug treatment and all experienced sterile protection, indicating that immunity from WM382-derived defective liver merozoites was long lasting (i.e. 10 months) and remained for 643 days following one challenge (**Fig. 1G, Table 1 Group 2**). Therefore, WM382 antimalarial activity against liver-stage merozoites not only provides chemoprophylaxis but results in durable immunogenicity that elicits sustained sterile protection against reinfection. This protective effect of an antimalarial PMIX/X dual inhibitor following a sporozoite infection is called CALM vaccination (**Fig. 1H**).

**Table 1.**
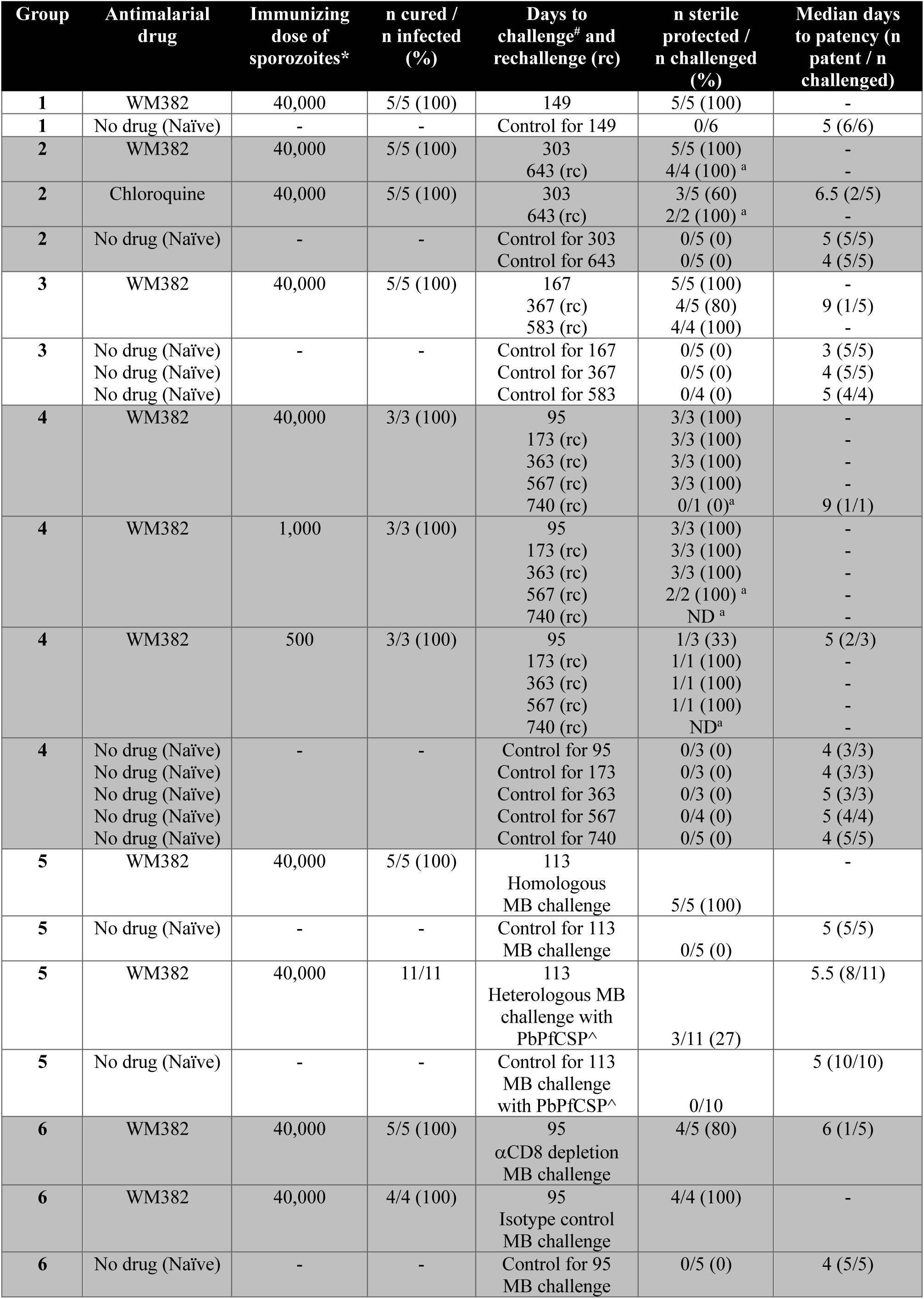
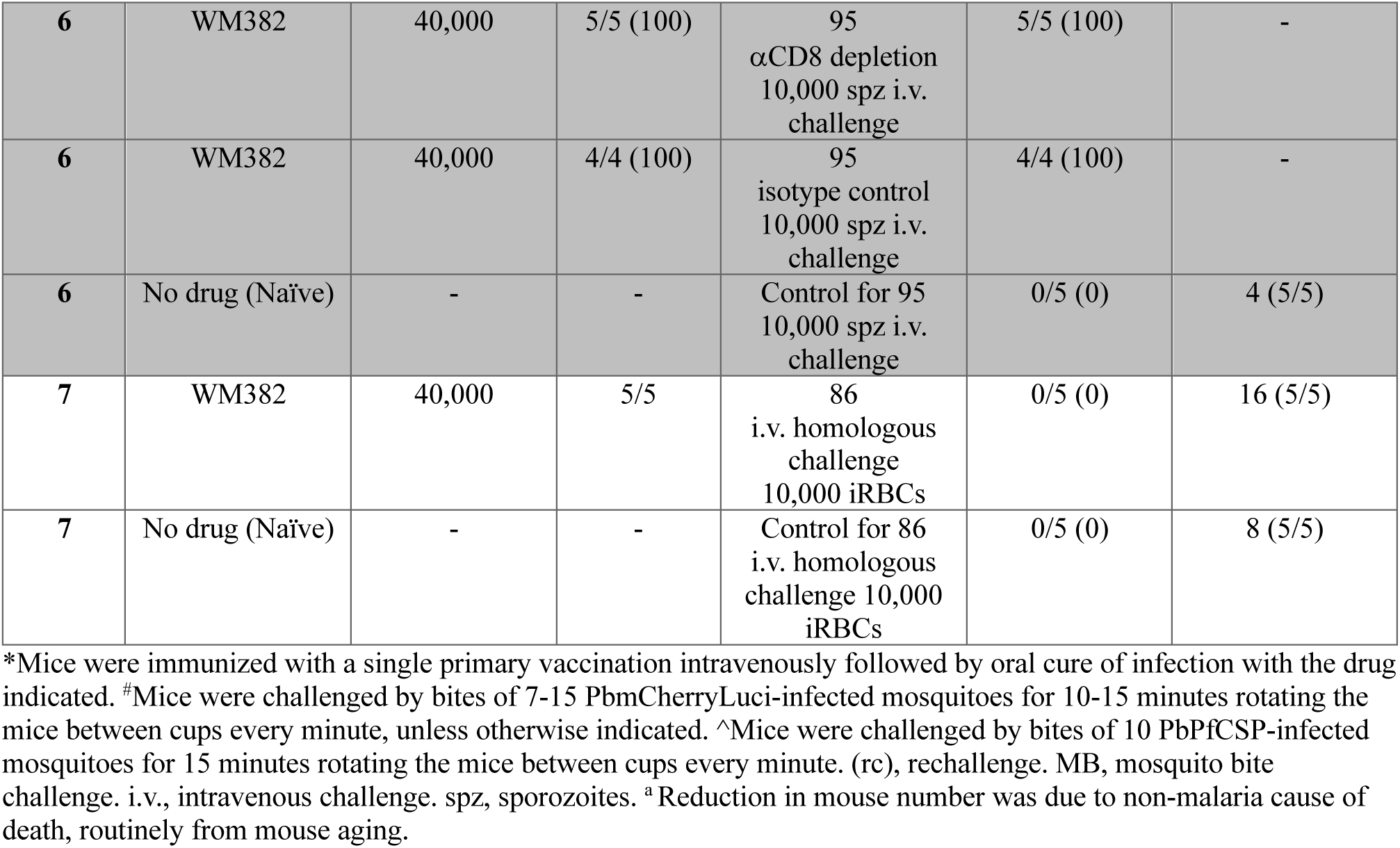
CALM vaccination of BALB/c mice.

| Group | Antimalarial drug | Immunizing dose of sporozoites* | n cured / n infected (%) | Days to challenge<sup>#</sup> and rechallenge (rc) | n sterile protected / n challenged (%) | Median days to patency (n patent / n challenged) |
| --- | --- | --- | --- | --- | --- | --- |
| **1** | WM382 | 40,000 | 5/5 (100) | 149 | 5/5 (100) | - |
| **1** | No drug (Naïve) | - | - | Control for 149 | 0/6 | 5 (6/6) |
| **2** | WM382 | 40,000 | 5/5 (100) | 303<br>643 (rc) | 5/5 (100)<br>4/4 (100)<sup>a</sup> | -<br>- |
| **2** | Chloroquine | 40,000 | 5/5 (100) | 303<br>643 (rc) | 3/5 (60)<br>2/2 (100)<sup>a</sup> | 6.5 (2/5)<br>- |
| **2** | No drug (Naïve) | - | - | Control for 303<br>Control for 643 | 0/5 (0)<br>0/5 (0) | 5 (5/5)<br>4 (5/5) |
| **3** | WM382 | 40,000 | 5/5 (100) | 167<br>367 (rc)<br>583 (rc) | 5/5 (100)<br>4/5 (80)<br>4/4 (100) | -<br>9 (1/5)<br>- |
| **3** | No drug (Naïve)<br>No drug (Naïve)<br>No drug (Naïve) | - | - | Control for 167<br>Control for 367<br>Control for 583 | 0/5 (0)<br>0/5 (0)<br>0/4 (0) | 3 (5/5)<br>4 (5/5)<br>5 (4/4) |
| **4** | WM382 | 40,000 | 3/3 (100) | 95<br>173 (rc)<br>363 (rc)<br>567 (rc)<br>740 (rc) | 3/3 (100)<br>3/3 (100)<br>3/3 (100)<br>3/3 (100)<br>0/1 (0)<sup>a</sup> | -<br>-<br>-<br>-<br>9 (1/1) |
| **4** | WM382 | 1,000 | 3/3 (100) | 95<br>173 (rc)<br>363 (rc)<br>567 (rc)<br>740 (rc) | 3/3 (100)<br>3/3 (100)<br>3/3 (100)<br>2/2 (100)<sup>a</sup><br>ND<sup>a</sup> | -<br>-<br>-<br>-<br>- |
| **4** | WM382 | 500 | 3/3 (100) | 95<br>173 (rc)<br>363 (rc)<br>567 (rc)<br>740 (rc) | 1/3 (33)<br>1/1 (100)<br>1/1 (100)<br>1/1 (100)<br>ND<sup>a</sup> | 5 (2/3)<br>-<br>-<br>-<br>- |
| **4** | No drug (Naïve)<br>No drug (Naïve)<br>No drug (Naïve)<br>No drug (Naïve)<br>No drug (Naïve) | - | - | Control for 95<br>Control for 173<br>Control for 363<br>Control for 567<br>Control for 740 | 0/3 (0)<br>0/3 (0)<br>0/3 (0)<br>0/4 (0)<br>0/5 (0) | 4 (3/3)<br>4 (3/3)<br>5 (3/3)<br>5 (4/4)<br>4 (5/5) |
| **5** | WM382 | 40,000 | 5/5 (100) | 113<br>Homologous MB challenge | 5/5 (100) | - |
| **5** | No drug (Naïve) | - | - | Control for 113 MB challenge | 0/5 (0) | 5 (5/5) |
| **5** | WM382 | 40,000 | 11/11 | 113<br>Heterologous MB challenge with PbPfCSP<sup>^</sup> | 3/11 (27) | 5.5 (8/11) |
| **5** | No drug (Naïve) | - | - | Control for 113 MB challenge with PbPfCSP<sup>^</sup> | 0/10 | 5 (10/10) |
| **6** | WM382 | 40,000 | 5/5 (100) | 95<br>αCD8 depletion MB challenge | 4/5 (80) | 6 (1/5) |
| **6** | WM382 | 40,000 | 4/4 (100) | 95<br>Isotype control MB challenge | 4/4 (100) | - |
| **6** | No drug (Naïve) | - | - | Control for 95 MB challenge | 0/5 (0) | 4 (5/5) |
| 6 | WM382 | 40,000 | 5/5 (100) | 95<br>αCD8 depletion<br>10,000 spz i.v.<br>challenge | 5/5 (100) | - |
| 6 | WM382 | 40,000 | 4/4 (100) | 95<br>isotype control<br>10,000 spz i.v.<br>challenge | 4/4 (100) | - |
| 6 | No drug (Naïve) | - | - | Control for 95<br>10,000 spz i.v.<br>challenge | 0/5 (0) | 4 (5/5) |
| 7 | WM382 | 40,000 | 5/5 | 86<br>i.v. homologous<br>challenge<br>10,000 iRBCs | 0/5 (0) | 16 (5/5) |
| 7 | No drug (Naïve) | - | - | Control for 86<br>i.v. homologous<br>challenge 10,000<br>iRBCs | 0/5 (0) | 8 (5/5) |

\*Mice were immunized with a single primary vaccination intravenously followed by oral cure of infection with the drug indicated. #Mice were challenged by bites of 7-15 PbmCherryLuci-infected mosquitoes for 10-15 minutes rotating the mice between cups every minute, unless otherwise indicated. ^Mice were challenged by bites of 10 PbPfCSP-infected mosquitoes for 15 minutes rotating the mice between cups every minute. (rc), rechallenge. MB, mosquito bite challenge. i.v., intravenous challenge. spz, sporozoites. <sup>a</sup> Reduction in mouse number was due to non-malaria cause of death, routinely from mouse aging.

Next, the efficacy of WM382 CALM was benchmarked against CQ chemoprophylaxis vaccination (CVac (*39*)), also known as chemoprophylaxis with sporozoites (CPS (*54*)) that elicits efficacious immunity in mice (*55*) and in clinical trials (*39, 54*). While CQ CVac induces potent antimalarial immunity, the emergence and spread of CQ resistance across the globe precludes the use of this drug in vaccination programs. CQ is distinct from WM382 in that it has no antimalarial effect on liver parasites (*42*), instead permitting complete liver stage development before exerting its activity on blood stage parasites following erythrocyte infection (*55*). Mice infected with 40,000 PbmChLuc sporozoites and treated with WM382 at the end of the liver stage or with CQ at the blood stage were challenged by mosquito bites 10 months (303 days) later. Both immunizations provided pre-erythrocytic immunity; however, virulent malaria breakthroughs occurred in some CQ CVac mice at this timepoint, with a median pre-patent period of 6.5 days compared with 5 days for naive controls, whereas all mice that received WM382 CALM retained sterile immunity (**Fig. 1F, G, Table 1 Group 2**). Surviving mice were rechallenged at 21 months (643 days) and all had sterile protection (**Fig. 1G, Table 1 Group 2**). Therefore, CALM conferred sterile immunity that was as efficacious and durable as the current gold-standard sporozoite vaccine CQ CVac, demonstrating the immunogenicity of liver-derived merozoites attenuated with WM382. As resistance to dual PMIX/X inhibitors is difficult to generate experimentally (*46, 47*), CALM represents a novel means to inhibit the parasite at the optimal late liver stage, generating a robust immunogenic mechanism of action for chemovaccination.

### CALM chemovaccination protects mice against repeated mosquito challenges for up to 2 years, including with low-dose immunizations

Humans living in malaria endemic areas are exposed to repeated infectious mosquito bites throughout the malaria transmission season, which typically lasts 3 to 6 months. Thus, to understand whether CALM would protect against repeated mosquito bite challenges, a primary vaccination with 40,000 sporozoites (**Fig. 1H**) was performed before exposing mice to multiple rounds of infectious mosquito bites over time without any further drug treatment. In the first experiment, mice that had previously received CALM and were fully protected when challenged at 5 months (167 days; **Fig. 1E**), were rechallenged at 12 months (367 days); four of the five mice were sterile protected, while the one breakthrough mouse experienced a 5-day delay to patency demonstrating significant pre-erythrocytic protection (**Fig. S1C, Table 1 Group 3**). A third challenge was given to the surviving mice at 19 months (583 days) and all retained sterile protection (**Fig. S1C, Table 1 Group 3)**. In the second experiment, CALM with 40,000 sporozoites conferred 100% sterile protection against four lethal challenges given at 3, 6, 12, and 19 months (95, 173, 363, and 567 days) and a 5-day delay to patency (but non-sterilizing immunity) when a fifth challenge was given at 24 months (740 days) (**Fig. S2F [40,000 SPZ]; Table 1 Group 4**). Therefore, WM382 CALM induces robust pre-erythrocytic immunity that protects against repeated challenges over extended time periods that were otherwise lethal to naive mice.

Others have shown that efficacy of sporozoite vaccines is related to the immunizing dose of sporozoites, with higher numbers of radiation-attenuated sporozoites (RAS) conferring greater efficacy than lower doses (*11*). However, as liver stage parasites mature, they divide exponentially and increase in size, biomass, and antigenic diversity, countering the need for very high sporozoite doses for efficacious immunity to be attained (*26*). Given the activity of WM382 is against late liver stage merozoites (after sporozoites entering the liver have replicated by many thousand-fold and developed to maturity), we measured the efficacy of CALM using lower immunizing doses of 500 and 1,000 sporozoites and compared this to 40,000 sporozoites (from the second experiment above) (**Fig. S2A, B**). While this cohort of nine vaccinated mice was not large, it showed consistently high levels of robust pre-erythrocytic immunity, including sterile protection against four challenges at 3, 6, 12 and 19 months in seven of the nine mice given a single CALM immunization with 500, 1,000 or 40,000 sporozoites, including all mice immunized with 1,000 sporozoites (*P*<0.0275; **Fig. S2C-F, Table 1 Group 4**). Taken together, our results show that a single WM382 CALM vaccination confers potent immunity that protects against multiple exposures to infected mosquito bites over lengthy time periods. Importantly, we provide evidence that immunizing doses ranging from 500 to 1,000 sporozoites provided robust and durable immunity. The potency of protection was likely due to the unique timing of WM382 against mature liver stage merozoites.

### CALM induces robust humoral responses including to CSP that neutralize sporozoites

Numerous studies have implicated the protective role of anti-CSP antibodies against sporozoites as they move from the dermal inoculation site to the liver (*56–58*). To assess whether CALM produces antibody responses and to assess if they are functional, several experiments were conducted. First, plasma from naive and vaccinated mice was collected before any challenge was given and tested for reactivity against *P. berghei* sporozoites and liver stages by immunofluorescence microscopy assay (IFA). Immune sera from mice vaccinated with CALM reacted with the periphery of both sporozoites and exoerythrocytic forms (**Fig. 2A,B**). As controls, immune sera from mice vaccinated with CVac and collected before challenge also reacted with the sporozoite periphery, as did anti-PbCSP 3D11 monoclonal antibodies, while pooled sera from naive mice was not cross-reactive (**Fig. 2A**). Second, to determine whether CSP was an antigen, IFA was performed on transgenic *P. berghei* sporozoites in which the native *CSP* gene was replaced with *P. falciparum CSP* (PbPfCSP) (*59*). Immune serum from neither CALM or CVac, nor anti-PbCSP 3D11 antibodies, cross-reacted with PbPfCSP sporozoites, whereas anti-PfCSP 2A10 monoclonal control antibodies labelled the sporozoite periphery but not PbmChLuc sporozoites as expected (**Fig. 2A**). This showed that immune sera from CALM reacted with PbCSP of sporozoites. Third, enzyme-linked immunosorbent assay (ELISA) analysis of immune serum taken before any challenges were given confirmed that CALM induced robust and sustained anti-PbCSP IgG responses in mouse serum lasting at least 19 months (581 days) (**Fig. 2C**). Anti-CSP titres were similar to those induced by CQ CVac when assessed 10 months (302 days) after vaccination (**Fig. 2D**). Fourth, to find whether anti-CSP humoral immunity from CALM was functional, immune sera from vaccinated mice that had not been challenged was incubated with PbmChLuc sporozoites dissected from mosquito salivary glands before the parasites were incubated with hepatocytes. Quantification of intracellular parasites 48 hours later revealed that CALM immune sera strongly inhibited hepatocyte infection by PbmChLuc sporozoites, similar to anti-PbCSP 3D11 control antibodies(*60*) and serum from CVac (**Fig. 2E, S3A, B**). CALM immune sera had no inhibitory effect on the infection of hepatocytes by PbPfCSP sporozoites, whereas an anti-PfCSP 2A10 monoclonal antibody control inhibited their invasion as expected (*61*) (**Fig. 2F, S3C**). Therefore, CALM vaccination induces sporozoite infection-blocking antibodies largely against CSP.

**Fig. 2.**
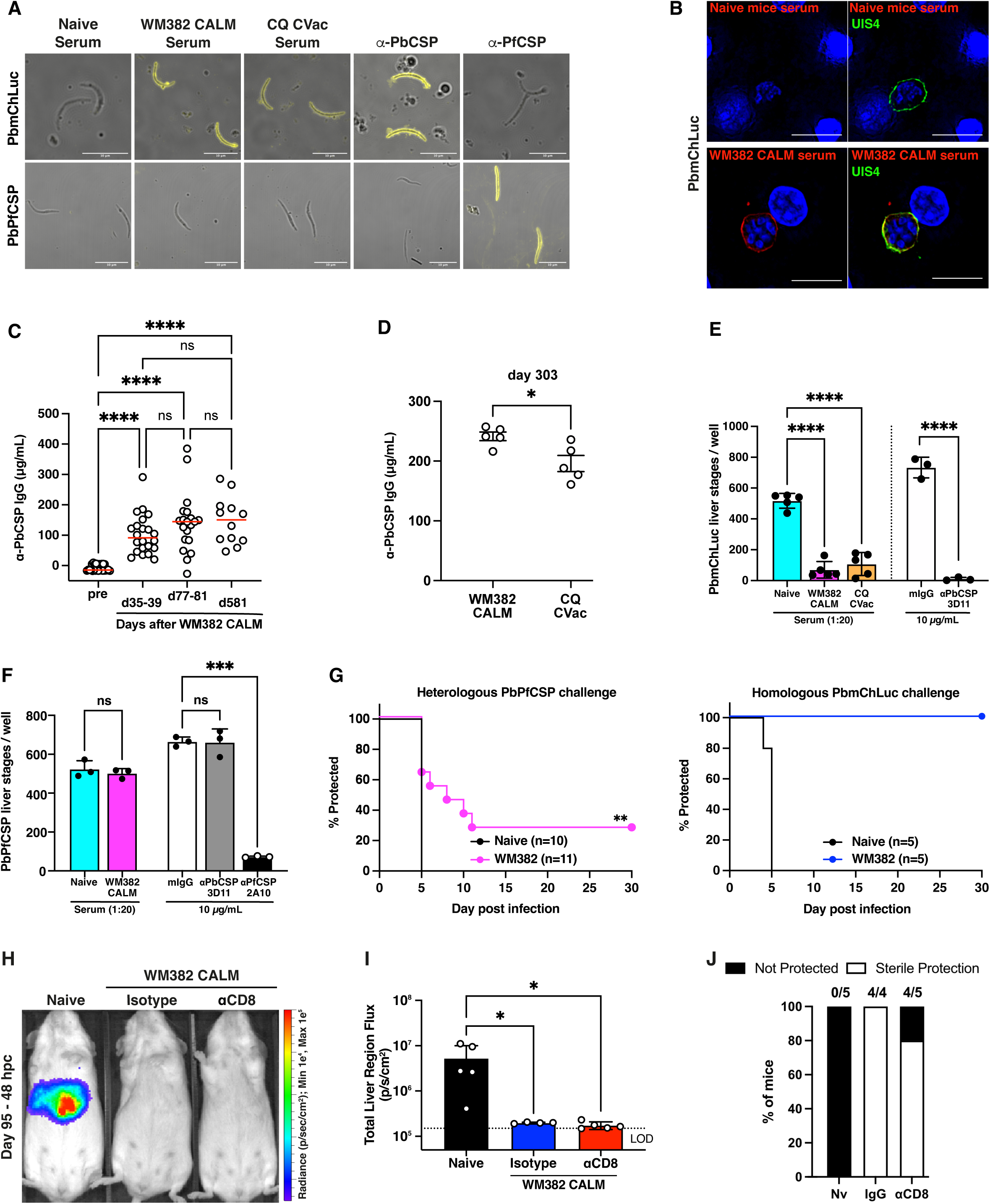
CALM vaccination stimulates humoral immunity in mice. **(A)** Immune serum from vaccinated mice (5 per group pooled, 1:100 dilution) recognizes *P. berghei* salivary gland sporozoites expressing mCherry-luciferase (PbmChLuc) but not *P. berghei* sporozoites expressing only CSP from *P. falciparum* (PbPfCSP). Scale bar, 10 µm. **(B)** Immune serum from vaccinated mice (5 per group pooled, 1:100) recognizes PbmChLuc liver stages within HepG2 cells fixed 25 hours postinfection (hpi). Scale bar, 10 μm. **(C)** Anti-PbCSP IgG titres measured by enzyme-linked immunosorbent assay (ELISA) at indicated days (D) after a single WM382 CALM vaccination of BALB/c mice or before CALM (pre). Data are pooled from one to three independent experiments (n=5-12 mice per experiment). Circles represent individual mice in technical replicates and median is shown (red). Data analyzed by one-way ANOVA with Tukey multiple comparisons test; ns, not significant; \*\*\*\**P*<0.0001. **(D)** Anti-PbCSP IgG titres measured by ELISA at day 303 after a single WM382 CALM or CQ CVac vaccination of BALB/c mice. Data are from one independent experiment (n=5 mice per group). Circles represent individual mice. Data analyzed by unpaired *t*-test with Welch correction; \**P*<0.05. **(E)** Number of PbmChLuc liver stages per well after incubation of sporozoites with immune serum from CALM or CVac mice (1:20 dilution) or control antibodies. Circles represent serum from individually vaccinated mice (left) and control antibodies (right) from 3 independent experiments. Data are mean ± SD analyzed by one-way ANOVA with Holm-Šídák multiple comparisons test or unpaired *t*-test (right); \*\*\*\**P*<0.0001. **(F)** Number of PbPfCSP liver stages per well after incubation of sporozoites with immune serum from CALM vaccinated mice or three control antibodies (right). Circles represent technical replicates for each condition. Mean ± SD. Data were analyzed by one-way ANOVA with Holm-Šídák multiple comparisons test or unpaired *t*-test. ns, not significant; \*\*\**P*<0.001. **(G)** WM382 CALM-vaccinated BALB/c mice were challenged 113 days later, together with naive controls, with bites from 10 PbPfCSP (pink) or PbmChLuc (blue) mosquitoes and monitored for blood stage parasites for 30 days. Data analyzed by log-ranked test; \*\**P*<0.01. **(H)** WM382 CALM vaccinated mice were IP treated with isotype control or anti-CD8 depleting antibodies 94 days later and challenged with bites of 10 PbmChLuc mosquitoes one day later. Representative IVIS imaging shows parasite liver loads 48 hours postchallenge (hpc). **(I)** Liver bioluminescence signals in all mice from (H) 48 hours postchallenge. Circles represent individual mice. Dashed line is mean signal of the pelvic region of all mice and the limit of detection (LOD). Mean ± SD. Data analyzed by one-way ANOVA with Holm-Šídák multiple comparisons test; \*\*\**P*< 0.001. **(J)** Percentage of mice from (I) with sterile protection (white) or not protected (black) against bites from10 PbmChLuc mosquitoes, measured by blood parasitemia for 30 days. Numbers above bars show the number of protected out of the total mice challenged, derived from one independent experiment (n=4-5 mice per group).

Finally, to demonstrate the importance of CALM-mediated anti-CSP immunity *in vivo*, vaccinated BALB/c mice were challenged 3 months (113 days) later with bites of mosquitoes infected with PbPfCSP or PbmChLuc control sporozoites. A total of 73% of mice challenged with heterologous PbPfCSP sporozoites succumbed to blood stage infection with a median prepatent period of 5.5 days, similar to naive controls, indicating significant loss of protection in those mice, in contrast to challenge with homologous sporozoites that resulted in complete sterile protection as expected (**Fig. 2G, Table 1 Group 5**). This indicated that anti-CSP immunity afforded by CALM vaccination conferred functional sterile protection *in vivo*. However, the remaining 27% of vaccinated mice were completely protected against PbPfCSP sporozoites, suggesting immunity was also occurring through other mechanisms. Altogether, these results indicate that attenuation of liver merozoites with WM382 elicited functional anti-CSP humoral immunity and suggests additional immune mechanisms may also contribute.

CD8^+^ T cells contribute to malaria liver stage protection in BALB/c mice by recognizing diverse antigens including an epitope in CSP (*62*). To test this possibility, CALM-vaccinated mice were depleted of CD8^+^ T cells (**Fig. S3D**) 1 day prior to their first challenge with homologous sporozoites by either mosquito bites or intravenous (IV) injection. Imaging with an *in vivo* imaging system (IVIS) revealed similarly low levels of liver parasite load between vaccinated mice treated with CD8-depleting or isotype control antibodies (**Fig. 2H, I; Fig. S3E, F**). In total, 80% and 100% of CD8-depleted mice maintained sterile protection against mosquito bites (**Fig. 2J**) and IV sporozoite challenge (**Fig. S3G**), respectively, with the one breakthrough mouse having a 2-day delay to patency (**Table 1 Group 6**). Together, these results indicate that WM382 CALM induces robust CSP-mediated humoral immunity against sporozoite challenges in BALB/c mice.

### PMIX/X inhibitors provide chemoprophylaxis and sterile immunity in B6 and outbred mice

Thus far our data demonstrated potency of WM382 against *P. berghei* liver stages in BALB/c mice (*46*), which generated durable sterile immunity (**Fig. 1, 2**). To extend these findings to a more stringent malaria mouse model that is susceptible to fatal disease without antimalarial intervention (*63, 64*), C57BL/6 (B6) mice were infected with 40,000 PbmChLuc sporozoites IV followed by oral administration of 100 mg/kg (mpk) WM382 at 36- and 48-hours postinfection (hpi) (**Fig. S4A**). IVIS quantification of parasite bioluminescence revealed normal liver stage development (52 hpi) and egress kinetics (55 hpi) in WM382-treated mice relative to no treatment controls (**Fig. S4B-D**). At 70 hpi, a systemic blood stage infection was observed by IVIS and flow cytometry in the untreated controls, but not in WM382-treated mice (**Fig. S4B, E-F**). Further examination demonstrated complete chemoprophylaxis of WM382-treated mice at the blood stage, manifest as the absence of parasitemia up to day 30 postinfection (**Fig. S4G**). These results were consistent with the dynamics of very late liver stage killing by WM382 in BALB/c mice that we reported previously (*46*). Therefore, WM382 given to B6 mice did not impede the development of liver stage parasites nor did it cause an obvious accumulation of liver parasites during their egress window; rather, it rendered liver merozoites defective and noninfectious as they were unable to initiate a blood-stage infection.

Having established comparable activity of WM382 against liver stage merozoites in both BALB/c and B6 mice, we next investigated the immunizing effect of drug treatment in the latter species. B6 mice were given a single dose of 40,000 sporozoites under WM382 cover dosed at 36 and 48 hpi to treat infection, representing a primary vaccination (CALM P), before being challenged with bites from 10 PbmChLuc infectious mosquitoes 4 months (108-133 days) later (**Fig. S5A**). Despite a significant reduction of liver parasite load (**Fig. S5B, C**), this single vaccination of B6 mice elicited suboptimal sterile protection of less than 10% (**Fig. S5D, Table 2 Groups 1-2**).

**Table 2.** CALM vaccination of C57BL/6 mice.

| Group | Antimalarial drug | Immunizing dose of sporozoites* | n cured / n infected (% cured) | Days to challenge<sup>#</sup> and rechallenge(rc) | n sterile protected / n challenged (% protected) | Median days to patency (n patent / n challenged) |
| --- | --- | --- | --- | --- | --- | --- |
| 1 | WM382 | 40,000 prime | 4/4(100) | 108 | 0/4 (0) | 5 (4/4) |
| 1 | MK-7602 | 40,000 prime | 2/2 (100) | 108 | 0/2 (0) | 4 (2/2) |
| 1 | No drug (Naïve) | - | - | Control for 108 | 0/3 | 4 (3/3) |
| 2 | WM382 | 40,000 prime | 6/6 (100) | 119 | 0/6 (0) | 4.5 (6/6) |
| 2 | WM382 | 40,000 prime | 6/6 (100) | 133<br>225 (rc)<br>365 (rc) | 1/6 (16.7)<br>1/1 (100)<br>0/1 (0) | 5 (6/6)<br>-<br>8 (1/1) |
| 2 | No drug (Naïve) | - | - | Control for 119/133<br>Control for 225<br>Control for 365 | 0/4 (0)<br>0/3 (0)<br>0/2 (0) | 4 (4/4)<br>3 (3/3)<br>4 (2/2) |
| 3 | WM382 | 40,000 prime<br>20,000 boost | 6/6 (100)<br>6/6 (100) | 113<br>Homologous challenge | 6/6 (100) | - |
| 3 | No drug (Naïve) | - | - | Control for 113<br>Homologous challenge | 0/6 (0) | 4 (6/6) |
| 3 | WM382 | 40,000 prime<br>20,000 boost | 6/6 (100)<br>6/6 (100) | 113<br>Heterologous PbPfCSP challenge<sup>^</sup> | 6/6 (100) | - |
| 3 | No drug (Naïve) | - | - | Control for 113<br>Heterologous PbPfCSP challenge<sup>^</sup> | 0/6 (0) | 4 (6/6) |
| 4 | WM382 | 40,000 prime | 4/4 (100) | 113<br>Isotype control i.v. 10,000 spz homologous challenge | 0/4 (0) | 5 (4/4) |
| 4 | WM382 | 40,000 prime | 4/4 (100) | 113<br>αCD8 depletion i.v. 10,000 spz homologous challenge | 0/4 (0) | 4 (4/4) |
| 4 | No drug (Naïve) | - | - | Control for 113 | 0/4 (0) | 3 (4/4) |
| 5 | WM382 | 40,000 prime<br>20,000 boost | 5/5 (100)<br>5/5 (100) | 149 | 4/5 (80) | 6 (1/5) |
| 5 | No drug (Naïve) | - | - | Control for 149 | 0/6 (0) | 4 (6/6) |
| 6 | WM382 | 40,000 prime<br>20,000 boost | 6/6 (100)<br>6/6 (100) | 128<br>Isotype control i.v. 10,000 spz homologous challenge<br>481 (rc MB) | 6/6 (100)<br>3/6 (50) | -<br>6 (3/6) |
| 6 | WM382 | 40,000 prime<br>20,000 boost | 4/4 (100)<br>4/4 (100) | 128<br>αCD8 depletion i.v. 10,000 spz homologous challenge | 0/4 (0) | 4 (4/4) |
| **6** | No drug (Naïve) | - | - | Control for 128<br>i.v. 10,000 spz<br>challenge<br>Control for 481<br>(MB) | 0/5 (0)<br>0/6 (0) | 4 (5/5)<br>3 (6/6) |
| **7** | MK-7602 | 40,000 prime<br>20,000 boost | 5/5 (100)<br>5/5 (100) | 90<br>383 (rc)<br>571 (rc) | 5/5 (100)<br>5/5 (100)<br>1/5 (20) | -<br>-<br>7 (4/5) |
| **7** | No drug (Naïve) | - | - | Control for 90<br>Control for 383<br>Control for 571 | 0/6 (0)<br>0/6 (0)<br>0/6 (0) | 3.5 (6/6)<br>4 (6/6)<br>3 (6/6) |
| **8** | WM382 | 40,000 prime<br>20,000 boost | 5/5<br>4/4 | 86<br>i.v. 10,000 iRBCs<br>homologous<br>challenge | 0/5 (0) | Median 15 days<br>to death (5/5) |
| **8** | No drug (Naïve) | - | - | Control for 86<br>i.v. 10,000 iRBCs<br>homologous<br>challenge | 0/5 (0) | Median 8 days<br>to death (5/5) |

\*Mice were immunized by prime-boost intravenous injections 6-7 weeks apart. #Mice were challenged at the indicated day post-primary vaccination immunization by bites of 10 PbmCherryLuci-infected mosquitoes for 10-15 minutes rotating the mice between cups every minute. ^Mice were challenged by bites of 10 PbPfcSP-infected mosquitoes for 15 minutes rotating the mice between cups every minute. (rc), rechallenge. MB, mosquito bite challenge. i.v., intravenous challenge. spz, sporozoites.

B6 mice given a malaria vaccine can develop stronger immunity after two or more immunizations, in part due to boosting of cellular responses (*38, 63, 64*). By repeating WM382 CALM vaccinations 6 weeks apart (**Fig. 3A**), representing a prime-boost regimen (CALM PB), 100% sterile protection was achieved, demonstrated by the lack of parasite liver burden following mosquito bite challenge and absence of parasitemia in all mice challenged either by mosquito bites (113-day challenge; **Fig. 3B-D**) or by IV injection of 10,000 PbmChLuc sporozoites (day 128 challenge; see Isotype **Fig. 5D-F; Table 2 Groups 3 and 6**) that was otherwise lethal to naive control B6 mice. Mice that survived the day 113 challenge were re-challenged 5 months (149 days) after primary vaccination by bites of 10 mosquitoes and sterile protection was achieved in 80% of mice (**Fig. 3B-D; Table 2 Group 5**). Mice that survived the day 149 challenge were rechallenged 16 months (481 days) after primary vaccination and 50% sterile protection was achieved (**Table 2 Group 6**). Together, these results show that the PMIX/X dual inhibitor WM382 has antimalarial activity against *P. berghei* liver stage merozoites in B6 mice and that prime-boost CALM vaccinations confer durable sterile protection against mosquito bite and high-dose IV challenges in this more stringent mouse model of lethal malaria.

**Fig. 3.**
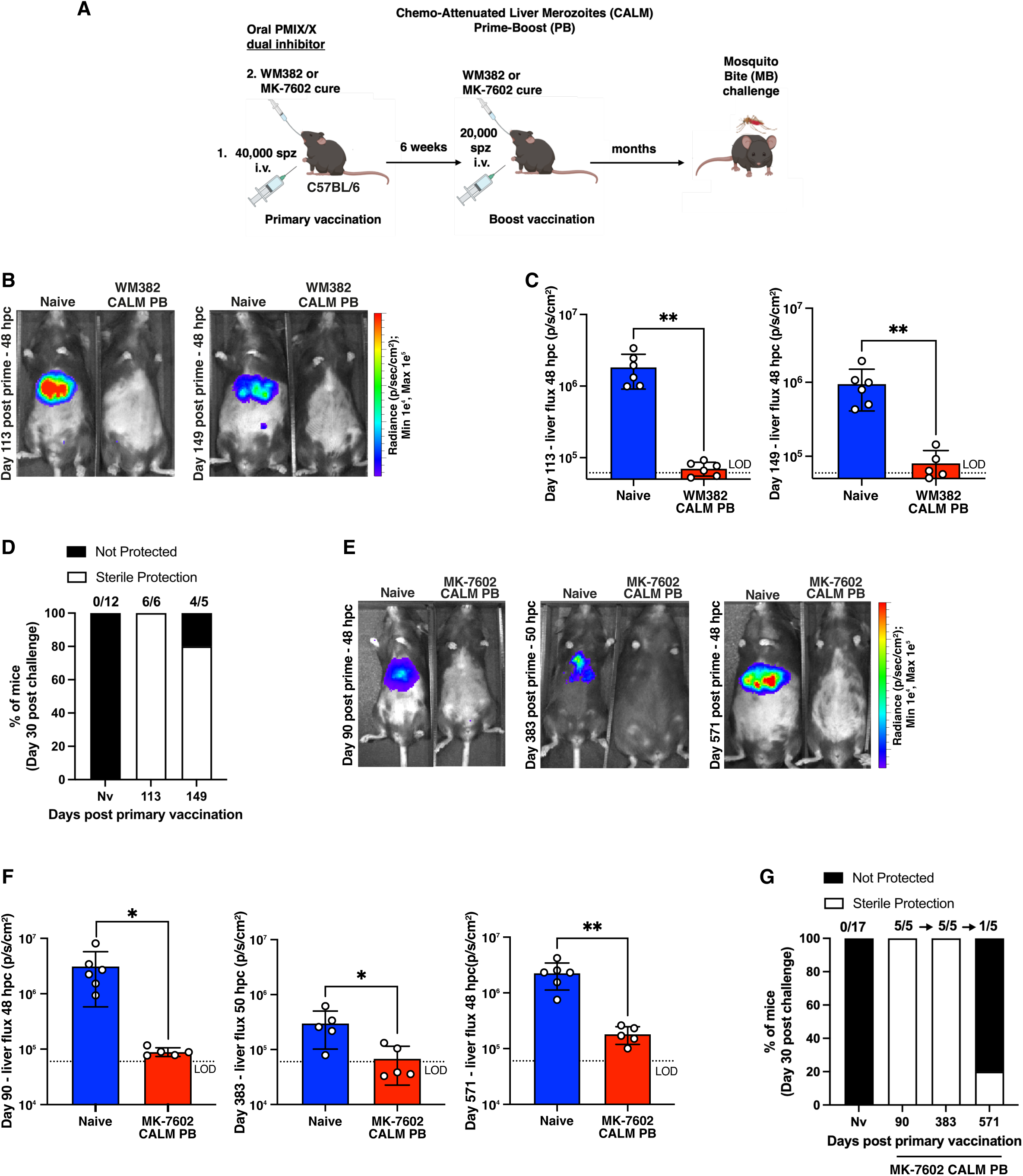
Prime-boost CALM vaccination protects C57BL/6 mice. **(A)** C57BL/6 mice given CALM vaccinations using WM382 or MK-7602 treatment at 36- and 48-hours following PbmChLuc sporozoite infection as a prime-boost (PB) regimen 6 weeks apart were challenged by bites of 10 PbmChLuc mosquitoes. **(B)** WM382 CALM PB vaccination protects mice against mosquito bite challenges at 113 and 149 days following primary vaccination, measured 48 hours postchallenge (hpc) by IVIS. **(C)** Liver bioluminescence signals in all mice from (B) 48 hours postchallenge (hpc). Circles represent individual mice. Dashed line is mean signal of the pelvic region of all mice and the limit of detection (LOD). Mean ± SD. Data analyzed by unpaired *t*-test; \*\**P*<0.01. **(D)** Percentage of mice from (C) with sterile protection (white) or not protected (black) against bites from 10 PbmChLuc mosquitoes, measured by blood parasitemia for 30 days. Numbers above bars show the number of protected out of the total mice challenged, derived from two independent challenge experiments. **(E)** MK-7602 CALM PB vaccination protects mice against mosquito bite challenges at 90 days, again at 383 days, and again at 571 days following primary vaccination, measured 48 to 50 hours postchallenge (hpc). **(F)** Liver bioluminescence signals in all mice from (E) 48 to 50 hours postchallenge (hpc). Circles represent individual mice. Dashed line is mean signal of the pelvic region of all mice and the limit of detection (LOD). Mean ± SD. Data analyzed by unpaired *t*-test; \**P*<0.05. **(G)** Percentage of mice from (F) with sterile protection (white) or not protected (black) against bites from 10 PbmChLuc infectious mosquitoes, measured by blood parasitemia for 30 days. Numbers above bars show the number of protected out of the total mice challenged, derived from one independent experiment (n=5-6 mice per group) in which surviving mice were rechallenged (for a total of three challenges).

To assess CALM vaccination in a highly stringent outbred model(*63, 64*), Swiss mice were infected with 40,000 sporozoites and treated with WM382 dosed at 36 and 48 hpi, confirming that 100% chemoprophylaxis could be achieved in this mouse species (**Fig. S6A**). As such, Swiss mice were given a primary vaccination using WM382 (CALM P) or CQ (CQ CVac P), the latter being a control with activity at the blood stage; 6 weeks later, half of the mice were given a booster immunization (CALM PB or CQ CVac PB) before all mice and naive controls were challenged by bites of 10 mosquitoes (**Fig. S6B, C**). Mice given a single vaccination and challenged 3 months later had reduced liver loads and developed blood stage infection with a 2-day delay to patency compared with naive mice, indicating immunity was generated by CALM and CQ CVac in Swiss mice but this was not infection blocking protection (**Fig. S6D-F; Table 3**). By contrast, prime-boost vaccination with CALM or CQ CVac achieved sterile immunity in 40% of mice, with the remaining mice experiencing 3-day and 2-day delays to patency, respectively, compared with naive controls (**Fig. S6D-F, Table 3**). Therefore, CALM vaccination elicits high levels of pre-erythrocytic and protective immunity in an outbred Swiss mouse model of lethal malaria with similar efficacy as the gold standard of CQ CVac (*36*).

**Table 3.** CALM vaccination of Swiss outbred mice.

| Group | Antimalarial drug | Immunizing dose of sporozoites* | n cured / n infected (% cured) | Days to challenge<sup>#</sup> | n sterile protected / n challenged (% protected) | Median days to patency (n patent / n challenged) |
| --- | --- | --- | --- | --- | --- | --- |
| 1 | WM382 | 40,000 prime | 4/4(100) | 83 | 0/4 (0) | 5.5 (4/4) |
| 1 | Chloroquine | 40,000 prime | 5/5 (100) | 83 | 0/5 (0) | 5 (5/5) |
| 1 | WM382 | 40,000 prime<br>20,000 boost | 5/5 (100)<br>5/5 (100) | 83 | 2/5 (40) | 6 (3/5) |
| 1 | Chloroquine | 40,000 prime<br>20,000 boost | 5/5 (100)<br>5/5 (100) | 83 | 2/5 (40) | 5 (3/5) |
| 1 | No drug (Naïve) | - | - | Control for 83 | 0/5 (0) | 3 (5/5) |

\*Mice were immunized by prime or prime-boost intravenous injections 6 weeks apart. <sup>#</sup>Mice were challenged at the indicated day post-primary vaccination immunization by bites of 10 PbmCherryLuci-infected mosquitoes for 15 minutes rotating the mice between cups every minute.

WM382 is a valuable lead compound that has elucidated novel PMIX and PMX biology across the *Plasmodium* life cycle (*46, 47*). To further validate that PMIX/X dual inhibition was effective in eliciting CALM vaccination, the clinical compound MK-7602 (*65*) was also evaluated. Our collaborative team developed this dual PMIX/X inhibitor with optimized drug-like properties and potential for clinical treatment of malaria (*65*). Like WM382, MK-7602 is active against liver-stage merozoites and completely attenuates the liver-to-blood transition following PbmChLuc sporozoite infection, providing chemoprophylaxis against malaria (*65*). To test whether MK-7602 elimination of liver merozoites was immunogenic like WM382 treatment, B6 mice immunized with MK-7602 CALM were evaluated for protection. As for WM382, an MK-7602 CALM primary vaccination afforded little protection (**Table 2, Group 1**); however, a boost vaccination conferred 100% sterile protection against 2 lethal challenges given 3 and 12 months (90 and 383 days) after primary vaccination and, after a third challenge at 19 months (571 days) 20% sterile protection was achieved with the susceptible mice experiencing 4-day delays to patency, indicating that immunity from MK-7602 treatment at immunization was durable (**Fig. 3E-G; Table 2 Group 7**). Therefore, CALM employing two different PMIX/X inhibitors with a novel late liver stage mechanism of action and a high barrier to resistance (*46, 65*), including the clinical drug MK-7602, induced robust pre-erythrocytic immunity that protected B6 mice against repeated mosquito bite challenges over extended periods that were otherwise lethal to naive B6 mice.

### CALM elicits mainly pre-erythrocytic rather than blood stage immunity

Liver merozoites express many blood stage proteins (*48*), raising the possibility that CALM could elicit protection against blood stage infection due to shared antigens. BALB/c mice were given a primary CALM vaccination (**Fig. 4A**), while B6 mice received prime-boost immunizations 6 weeks apart (**Fig. 4B**), before all mice and naive controls were challenged with 10,000 PbmChLuc-infected erythrocytes IV. While naive mice of both strains developed virulent malaria and required euthanasia between days 8 and 11 postinfection, CALM-vaccinated mice controlled their malaria symptoms and parasitemia for a median 8 days longer than naive controls (*P*<0.005) before requiring humane endpoint intervention due to clinical malaria (**Fig. 4C-F; Table 1 Group 7; Table 2 Group 8**). No further investigation was performed as to how this partial stage-transcending clinical immunity by CALM is mediated since it was not sterilizing. Rather, we focused on better understanding the cellular immunity that is protective against pre-erythrocytic infection following sporozoite challenge.

**Fig. 4.**
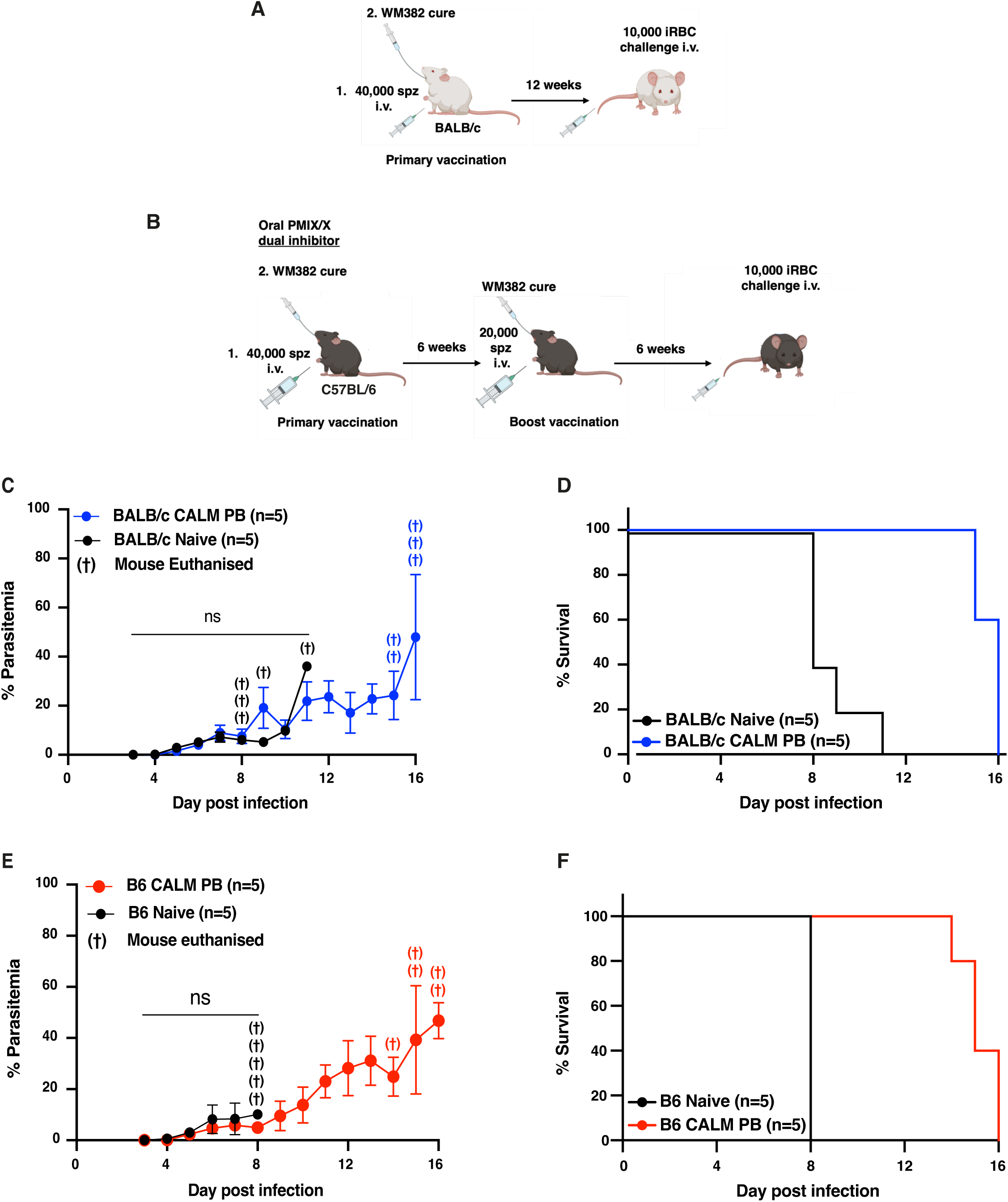
Stage transcending immunity by WM382 CALM. **(A)** WM382 CALM vaccinated BALB/c mice were challenged 12 weeks later IV with 10,000 PbmChLuc-infected erythrocytes and parasitemia monitored by microscopy for 16 days. **(B)** WM382 CALM prime-boost vaccinated C57BL/6 mice were challenged 6 weeks later IV with 10,000 PbmChLuc-infected erythrocytes and parasitemia monitored by microscopy for 16 days. **(C)** Parasitemia in WM382 CALM vaccinated from (A) and naive BALB/c mice. Humane endpoints requiring euthanasia are indicated (†). **(D)** Survival curves of BALB/c mice from (C) showing time to humane endpoint requiring euthanasia. **(E)** Parasitemia in WM382 CALM vaccinated from (B) and naive C57BL/6 mice. Humane endpoints requiring euthanasia are indicated. **(F)** Survival curves of C57BL/6 mice from (E) showing time to humane endpoint requiring euthanasia.

### CALM induces functional CD8^+^ T-cell immunity including long-lived Trm cells against diverse antigens

Our BALB/c mouse data revealed functional anti-CSP humoral immunity following a single immunization with WM382 CALM. To characterise this response in B6 mice, animals were provided a prime or prime-boost immunization and serum was analyzed by ELISA. This revealed that some, but not all, mice developed a small but significant increase in PbCSP IgG titres following vaccinations (**Fig. 5A**). B6 mice immunized via prime-boost with PbmChLuc sporozoites enjoyed 100% sterile protection against bites of 10 PbPfCSP infectious mosquitoes (**Fig. 5B, Table 2 Group 3**), suggesting that B6 mice were less dependent on humoral immunity to CSP than BALB/c mice, implicating alternate immune mechanisms.

**Fig. 5.**
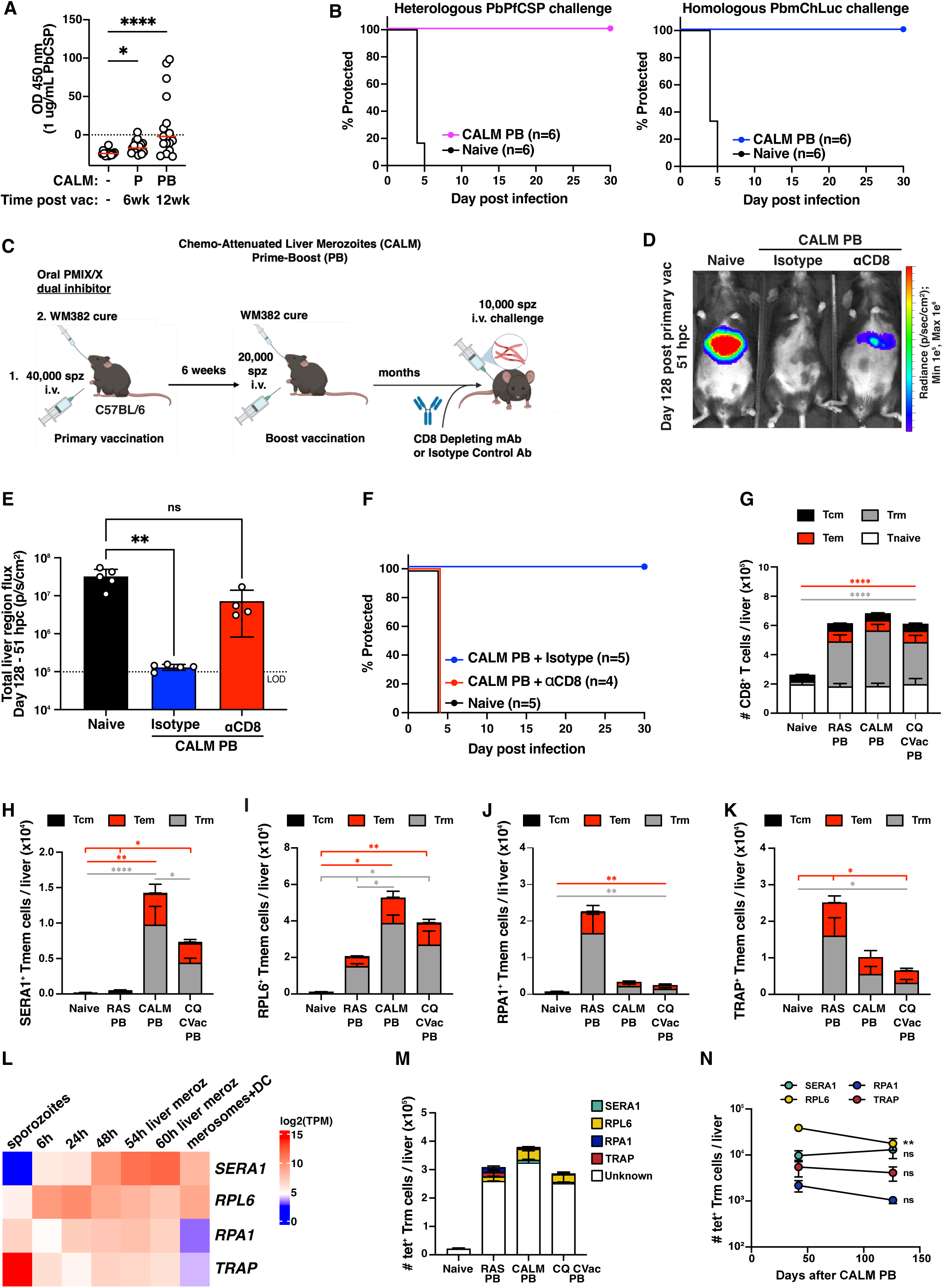
CALM induces humoral and CD8^+^ T cell immunity towards diverse antigens. **(A)** Anti-PbCSP IgG titres measured by ELISA 6 and 23 weeks (wk) after a primary (P) or prime-boost (PB) WM382 CALM vaccination regimen in B6 mice. Data are pooled from two independent experiments (n=17 mice). Circles represent individual mice and median is shown (red). Data analyzed by one-way ANOVA with Dunn’s multiple comparisons test; \**P*<0.05, \*\*\*\**P*<0.0001. **(B)** WM382 CALM prime-boost (PB)-vaccinated B6 mice were challenged 113 days later, together with naive controls, with bites from 10 PbPfCSP (pink) or PbmChLuc (blue) mosquitoes and monitored for blood stage parasites for 30 days. **(C)** WM382 CALM prime-boost (PB)-vaccinated B6 mice were IP treated with isotype control or anti-CD8 depleting antibodies once at 127 days following primary vaccination and challenged one day later by IV injection with 10,000 PbmChLuc sporozoites. **(D)** IVIS imaging of mice in (C) shows parasite liver loads 51 h postchallenge (hpc). **(E)** Liver bioluminescence signals in all mice from (D) at 51 hours postchallenge (hpc). Circles represent individual mice. Dashed line is mean signal of the pelvic region of all mice and the limit of detection (LOD). Mean ± SD. Data were analyzed by one-way ANOVA with Dunnett multiple comparisons test. ns, not significant; \*\**P*<0.01. **(F)** Survival curves show percentage of mice from (E) protected against IV challenge with 10,000 PbmChLuc sporozoites (n=4-5 mice per group) after 30 days. **(G)** Numbers of CD8^+^ T cells in livers of B6 mice immunized PB with either radiation-attenuated sporozoites (RAS), WM382 CALM, or CQ CVac quantified by flow cytometry 42 days after the boost immunization. Naive (Tnaive; CD44^-^ CD8^+^), tissue resident memory (Trm; CD44^+^ CD69^+^ CD62L^-^), effector memory (Tem; CD44^+^ CD69^-^ CD62L^-^), and central memory (Tcm; CD44^+^ CD69^-^ CD62L^+^) T cells shown. Counts are mean ± SEM compared by one-way ANOVA with Tukey’s multiple comparisons test; \*\*\*\**P*<0.0001 compared to Naive. **(H)** Number of tetramer-specific Trm cells in the liver 42 days following boost immunization specific for *P. berghei* SERA1 **(I)**, RPL6 **(J)** RPA1, **(K)**, and TRAP were quantified by flow cytometry. **(L)** RNAseq heatmap of *P. berghei* liver stage expression for antigens *SERA1* (serine repeat antigen 1), *RPL6* (60S ribosomal protein L6), *RPA1* (replication protein A1), and *TRAP* (thrombospondin-related anonymous protein) sampled from sporozoites through to merosomes (and detached cells; DCs) *in vitro*. Data mined from reference (*49*); h, hours postinfection; TPM, transcripts per million. **(M)** Total numbers of liver Trm cells specific to known (SERA1, RPL6, RPA1 and TRAP) and unknown *P. berghei* antigens 42 days following boost immunization. **(N)** Number of tetramer-specific Trm cells in the livers of B6 mice 42- and 126-days following boost immunization with WM382 CALM. Data in (H-K, M, N) are mean ± SEM pooled from two independent experiments (n=6-10 mice/group) compared by one-way ANOVA with Tukey multiple comparisons test; \*\**P*<0.01, \*\*\**P*<0.001, \*\*\*\**P*<0.0001. Data in **(N)** were log-transformed and compared by unpaired *t*-test; ns, not significant; \*\**P*<0.01.

To explore the protective role of cellular immunity, B6 mice treated with prime or prime-boost immunizations were depleted of CD8^+^ T cells 1 day prior to IV challenge with a high dose of 10,000 PbmChLuc sporozoites, while control-vaccinated mice received an isotype antibody that did not affect their CD8^+^ T cells (**Fig. 5C, S7A**). Since sporozoites can be neutralized by antibody responses in the skin and *enroute* to the liver (*57, 58*), this route was bypassed by using IV injection of sporozoites, which results in rapid invasion of hepatocytes (*66*). While B6 mice given a WM382 CALM primary vaccination and isotype antibodies experienced ∼85% reduction in parasite liver load (*P*=0.0134) and a median 2-day delay to patency compared with naive mice (*P*=0.0058), depletion of CD8^+^ T cells following CALM primary vaccination (**Fig. S7B**) resulted in little protection, manifest as liver loads that were similar to naive controls (*P*=0.6134) and a modest delay to patency (*P*=0.1859 Mantel-Cox Log-rank test) (**Fig. S7C-E**). Importantly, mice immunized with prime-boost vaccinations and depleted of their CD8^+^ T cells were susceptible to liver infection (**Fig. 5E**; *P*=0.2979 versus naive mice) and shared the same prepatent period as naive mice, while vaccinated mice that received an isotype antibody control retained liver stage immunity and were malaria free for 30 days, indicating they had sterile protection against a high dose challenge of 10,000 sporozoites IV (**Fig. 5D-F**). Having demonstrated the important function of CD8^+^ T cells in protection that was enhanced upon boosting, we undertook flow cytometry analyses of the CD8 compartments. This revealed the induction of memory CD8^+^ T cells, predominantly tissue resident memory (Trm) cells, in the livers of mice given WM382 CALM vaccinations that was substantially increased following the boost vaccination (**Fig. 5G, S5E**). Altogether, these results indicate that CD8^+^ T cells are induced, boosted, and required for sterile protection that was elicited by WM382 CALM vaccination in B6 mice.

Given the critical role of liver Trm cells in mediating liver stage immunity (*32*), this memory subset became the focus of our next analyses. Using MHC-I tetramers specific to four known malaria antigens (*67–70*), distinct compositions of SERA1, RPL6, RPA1 and TRAP-specific responses were identified following WM382 CALM. As this vaccine involves very late liver stage arrest, we also compared the responses of Trm cells induced by vaccination either with RAS, which involves early liver stage arrest, and CQ CVac that involves blood-stage killing after sporozoite infection. RAS and CQ CVac had comparable expansion of CD8^+^ T cells to CALM (**Fig. 5G, S5E**). Consistent with WM382 CALM being a liver merozoite-arresting vaccine, the memory CD8^+^ T cell responses were somewhat distinct from those to RAS and similar to the CQ CVac response (**Fig. 5H-K**), including generation of Trm cells reactive to *P. berghei* SERA1 and RPL6 that are strongly transcribed in liver merozoites and merosomes (**Fig. 5L)** and are protective antigens (*67, 70*). Irrespective of the vaccination strategy, a significant fraction of liver Trm cells exhibited antigen specificities that are under investigation but remain to be identified (**Fig. 5M**).

Lastly, a longitudinal examination showed that liver Trm cell populations of diverse antigen specificity remained relatively stable for at least 4 months (126 days) after the WM382 CALM boost immunization, although a small attrition in the number of RPL6-specific Trm cells was noted during this period (**Fig. 5N**). Collectively, these results demonstrate that CD8^+^ T cells are important for protection using CALM and that prime-boost immunization induces large and antigenically diverse pools of long-lived liver Trm cells.

### WM382 attenuates *P. falciparum* liver-stage merozoites in FRG huHep mice

To test the activity of PMIX/X dual inhibitors against the human malaria parasite at the liver stage, humanized FRG mice deficient of fumarylacetoacetate hydrolase (*<u>F</u>ah^-/-^*), immunocompromized (*<u>R</u>ag2^-/-^*, *Il2r<u>g</u>^-/-^*), and engrafted with human hepatocytes (huHep) were used. In this model, *P. falciparum* liver stage development takes 7 days, with parasites egressing on the last day (*71*). To this end, FRG huHep mice were intravenously infected with 300,000 *P. falciparum* NF54 sporozoites expressing GFP-luciferase (*72*), followed by oral dosing with WM382 on days 5 and 6 postinfection when liver merozoites are developing (**Fig. 6A**). IVIS imaging demonstrated comparable profiles of liver stage development in WM382-treated mice versus no treatment control mice up to day 6.8 postinfection (164 hpi) (**Fig. 6B, C**). As attenuation of *P. berghei* by WM382 prevents the first wave of erythrocytic infection (*46*) (and this study; **Figs. 1C, S4, S6A**), we assessed the liver-to-blood transition of *P. falciparum* by engrafting uninfected human erythrocytes into the liver-infected FRG huHep mice on day 6 and 6.5 postinfection, providing the red blood cell niche for egressing liver stage merozoites to invade and develop (**Fig. 6A, D, E**). Whole-body bioluminescence from no treatment control mice detected the first wave of *P. falciparum* blood stage infection on days 7 and 8 as expected, evidenced by bioluminescence emanating from parasites located in the tail, paws, head and chest from circulating blood, yet in WM382-treated mice, this distribution was restricted to the liver and very faintly in the chest where we cannot exclude the possibility that attenuated merozoites may have accumulated and/or died (**Fig. 6B**). Blood from all six FRG mice was collected on day 7.7 (184 hpi) and cultured *in vitro* with fresh human erythrocytes to allow parasite expansion. As expected, asexual blood stage parasites were identified in cultures from all three no treatment control mice 3 days later (day 11), and again 5 days later (day 16), indicating successful transition of *P. falciparum* liver stage merozoites into the blood phase of infection had occurred in each of those FRG mice on days 7 and 8 (**Fig. 6B**). By contrast, no blood stage parasites were observed in the *in vitro* cultures containing blood from all WM382-treated FRG mice by days 11 or 16, indicating that WM382 had greatly attenuated *P. falciparum* liver merozoites in the three FRG mice (**Fig. 6F**). A very small number of viable parasites may have been undetected during this culture period, so cultures from one of the WM382-treated mice was maintained *in vitro* for 8 weeks (out to day 64) and remained *P. falciparum* negative. This established that liver-to-blood transition was definitively prevented in this animal; this 8-week assessment was not possible in the remaining two WM382-treated animals due to their contamination after day 16 (**Fig. 6F**). Nonetheless, these results demonstrate that WM382 permits normal *P. falciparum* liver stage development up to day 6, and egress from the liver between days 6 to 7.7, like no treatment control parasites. However, WM382 attenuated the ability for *P. falciparum* to establish the first erythrocytic wave of infection for at least 16 days in all mice, including completely protecting at least one mouse for 30 days.

**Fig. 6.**
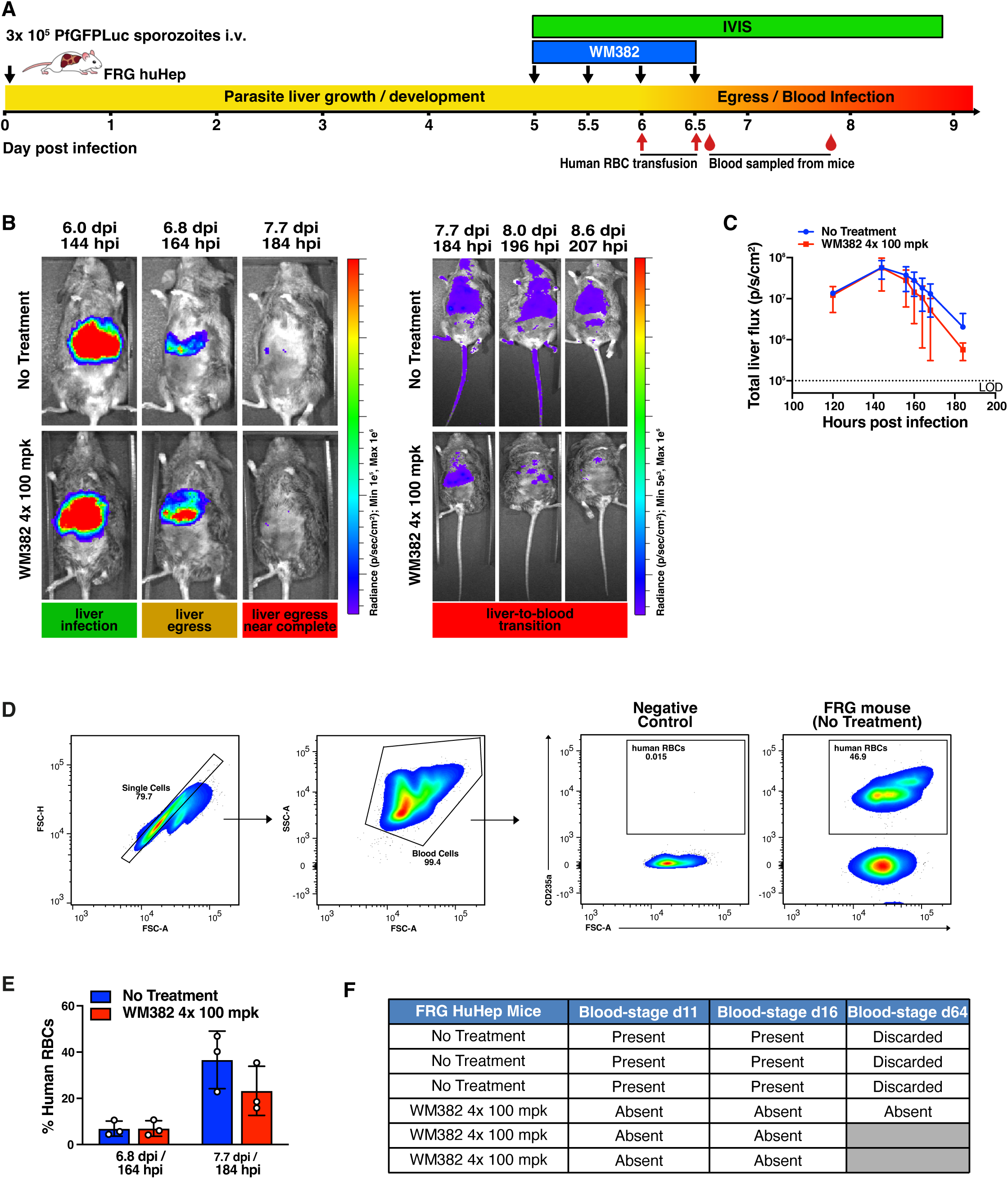
WM382 chemo-attenuation of *P. falciparum* liver-to-blood transition in FRG huHep mice. **(A)** Experiments for (b-f). Six FRG huHep mice infected IV with 300,000 *P. falciparum* NF54 sporozoites expressing GFP-Luciferase (PfGFPLuc) per mouse were treated with either WM382 (4x 100 mg/kg [mpk] orally 12 hours apart) from day 5 to 6.5 postinfection (n=3 mice) or No Treatment (n=3 mice). Liver infection was measured as PfGFPLuci bioluminescence by IVIS. FRG mice were transfused with human red blood cells (RBC) at day 6 and 6.5 with whole blood sampled at day 6.8 and 7.7 to confirm engraftment. **(B)** IVIS imaging of FRG huHep mice shows PfGFPLuci liver stage development and egress (left) and liver-to-blood transition using longer exposures (right) in the presence or absence of WM382. **(C)** PfGFPLuci bioluminescence signal of liver stage development and egress in all six FRG huHep mice imaged by IVIS. Dashed line is mean liver signal of all mice before luciferin injection and the limit of detection (LOD). Mean ± SD. **(D)** Representative FACS plots after transfusion of human red blood cells (RBC) into FRG huHep mice on day 6 and 6.5 postinfection, measuring engraftment in whole blood on days 6.8 and 7.7. **(E)** Percent engraftment of human RBCs in all six FRG huHep mice, measured by flow cytometry of whole blood samples collected 6.8- and 7.7-days postinfection (dpi) (164- and 184-h postinfection, hpi). Circles represent individual mice. Mean ± SD. **(F)** Blood sampled on day 7.7 from all six FRG huHep mice was cultured *ex vivo* with fresh human erythrocytes for up to 8 weeks (56 days or 64 d postinfection); d, days. Cultures were monitored for blood stage parasites by Giemsa thin blood smears and microscopy. Cultures from no treatment mice were all parasite-positive on day 11 after sporozoite infection, expanded exponentially and were discarded after day 16. Cultures from WM382 mice were maintained for 56 days, two samples developed yeast contamination from day 17 (greyed out).

## Discussion

Here, we report the unique property of first-in-class antimalarials, including the clinical compound MK-7602 (*46*), that completely attenuate the liver-to-blood transition of the malaria parasite *P. falciparum* and *P. berghei* through the dual inhibition of PMIX and PMX (*46*). This elicits durable sterile immunity in mice with similar efficacy to CPS/CQ CVac, representing a promising development. Attenuated sporozoite vaccines stimulate robust humoral and cellular responses including CD8^+^ cells that provide durable protection against reinfection. Although experimental immunization with sporozoites under CPS/CQ CVac provided up to 100% efficacy (*36, 39–41, 54, 73*), broad uptake was limited by global parasite resistance to CQ. Alternative antimalarials have been evaluated in chemovaccination studies, however, most attenuate parasites at suboptimal phases of liver or blood stage infection for optimal immune efficacy (reviewed in reference (*21*)). The late liver stage has been identified as an ideal inflection point to attenuate the parasite for broad spectrum, efficacious immune education to liver- and blood-stage antigens before blood-stage infection takes hold and erodes immunity (*22–25, 45, 74*). Very few drugs approved for malaria target this phase of the life cycle (*46, 50, 51, 75*). The recent discovery and clinical development of PMIX/X dual targeting antimalarials with a high barrier to resistance and potent activity against the very late liver stage (*46, 47, 65*) offers exciting potential.

The aspartyl proteases PMIX and PMX process many parasite substrates required for egress and invasion of host cells (*46, 50, 51, 76, 77*). Addition of PMIX/X inhibitors during hepatocyte infection permits the exponential replication and maturation phases of exoerythrocytic forms and results in only a modest reduction in merosome formation *in vitro*, but completely inhibits erythrocyte infection by non-viable, liver-derived merozoites (*46, 65*). This phenotype is likely due to inhibition of cleavage of numerous substrates (*48, 78, 79*), including multi-member protein families, that are required both for egress from hepatocytes and to initiate the first wave of erythrocytic infection. The phase of parasite arrest following WM382 / MK-7602 treatment of liver infection *in vivo* (*46, 65*) may be slightly later than reported for GA2 and LARC2 vaccines that arrest at liver schizonts (*23, 24*). Whether release of noninfectious merosomes from the liver is advantageous for priming functional immunity is not yet clear, but merosomes can lodge and rupture in the lungs (*4*), which were recently shown to be an important site for CD8^+^ T cell activation following sporozoite vaccination (*80*). Death of liver merozoites following WM382 / MK-7602 treatment also occurs inside hepatocytes (*46*) and this may also contribute to immune education *in vivo*.

PMIX and PMX are clearly important enzymes and drug targets in exoerythrocytic and blood stage merozoites. These proteases are also expressed in sporozoites isolated from both the oocysts and salivary glands of the mosquito (*49, 81, 82*). Pretreatment of mice with WM382 4 hours before infection with bites of *P. berghei*-infected mosquitoes had no deleterious effect on subsequent liver infection kinetics (*46*) suggesting that the function of these proteases in sporozoites may be related to biology within the mosquito. PMIX and PMX are not transcribed appreciably (*49*), nor are they essential (*46, 50*), in the early- or mid-phases of parasite development in hepatocytes (*46, 65*), possibly explaining why our compounds permit development to maturity of *P. berghei* and *P. falciparum* liver stages.

That PMIX/X inhibitors permit complete liver stage development before attenuating mature liver merozoites is salient in the context of malaria vaccines. Early liver stage arresting sporozoite vaccines require high doses to achieve sterile immunity (eg, 10^5^ to 10^6^ sporozoites or a minimum of 1,000 bites by irradiated mosquitoes) (*11–15, 41*). By contrast, late liver-stage-arresting vaccines (*22–24*) require fewer sporozoites and provide superior efficacy to those that arrest earlier (*11–15, 41*). The reasons are multifactorial and include the exponential growth of sporozoites into large parasites with augmented biomass and broad antigen expression that elicit strong CD8^+^ T cell responses including liver-resident Trm cells to diverse pre-erythrocytic antigens (*83*). The late liver stage is currently being pursued as an attractive immune target using genetically attenuated vaccines with exciting results (*23–26*). The late liver-stage attenuated *P. falciparum* sporozoite vaccine GA2 achieved ∼90% protection in volunteers that received one or three immunizations from the bites of 50 mosquitoes infected with GA2 (*25, 26*), directly demonstrating the potent immunogenicity of late liver stage attenuation. However, challenges remain in scale-up, commercial manufacture and necessity for IV injections. A chemotherapeutic approach to attenuating the late liver stage for vaccination against malaria offers some advantages, discussed below. Notably, this approach also comes with inherent risks, including the requirement for infection by wildtype sporozoites that can cause severe disease if drug resistance develops and/or breakthrough infections occur. The barrier to resistance for dual PMIX/X-specific antimalarials is presently high (*46, 47, 65*) and may be protected by targeting the liver-stage bottleneck where fewer parasites are exposed to drug than during blood-stage infection.

CALM chemovaccination provided sustained sterile immunity against repeated lethal sporozoite challenges in three mouse species for sustained periods. Immunity was predominantly pre-erythrocytic, although partial clinical immunity against blood stage parasites was conferred, likely due to attenuated liver-derived merozoites presenting both liver- and blood stage antigens. This contrasts with CQ CVac/CPS, which provided sterile pre-erythrocytic immunity but no protection against blood stage challenge (*84*).

The benefit of eliciting CD8^+^ T cell responses for immune protection is now well established (*28–38*). CALM vaccination stimulated robust CD8^+^ T cell responses, including Trm reactivity to diverse antigens including RPL6 (*67*) and the recently discovered protective antigen SERA1 (*70*), both of which are transcribed at high levels in liver merozoites and merosomes (*49*). SERA1 possesses a putative consensus motif for cleavage by PMIX and/or PMX, suggesting it may be yet another a substrate. CALM also stimulated robust anti-CSP antibody responses that neutralized sporozoites, providing a high degree of immune protection against future infection by mosquito bites and IV challenge. Therefore, CALM represents a chemotherapeutic means to achieve an outcome that is similar to late-arresting GAPs, using a novel class of antimalarials including MK-7602 that is now in clinical development (*65*). We showed that MK-7602 CALM provided durable sterile immunity in a stringent B6 mouse malaria model lasting for more than 1 year. *Plasmodium* species including *P. vivax* and *P. knowlesi* are, like *P. falciparum* and *P*. *berghei*, sensitive to PMIX/X inhibitors (*46, 65*) raising the possibility that CALM can be used to vaccinate against all circulating species and strains, which is highly desirable for malaria elimination.

CALM chemovaccination demonstrated high efficacy in mice, even with low sporozoite doses (10² to 10³), likely due to exponential replication of parasites and arrest at a very late stage. This has implications regarding the number of sporozoites necessary to induce functional immunity in humans under PMIX/X drug coverage and whether natural, repeated exposure to sporozoites under such drug coverage could progressively generate immunity, potentially obviating the need for direct sporozoite inoculation in humans (*20*). Sporozoite expulsion by *Anopheles* mosquitoes infected with both laboratory and naturally circulating *P. falciparum* gametocytes can result in surprisingly high inocula, averaging over 1,000 sporozoites per bite, and transmission is more likely in mosquitoes with higher salivary gland loads (*85, 86*). Remarkably, as few as three ruptured oocysts can generate salivary gland loads exceeding 10,000 *P. falciparum* sporozoites in experimentally infected *An. stephensi*, underscoring the efficiency with which this parasite leverages the mosquito vector to amplify its transmission potential (*86*). This efficiency suggests that even low-level infections may have significance in the context of natural chemovaccination, particularly in regions with high mosquito densities or frequent human-vector contact. In Burkina Faso, for instance, biting rates have been reported to range from 2.2 to 52.2 bites per person per night during the malaria season (*87*), highlighting the potential for repeated exposure. Collectively, these findings provide support for the notion that natural, repeated exposure to sporozoites while under PMIX/X-targeted drug coverage could progressively drive the development of protective immunity.

A promising strategy for translating CALM to endemic regions involves the development of long-acting injectable (LAI) compounds and appropriate formulations of PMIX/X inhibitors. The example of lenacapavir as HIV prophylaxis, providing up to 6 months of protection with a single injection, suggests such a pathway (*88*). A once-per-season LAI could become a transformative tool in malaria elimination. An LAI targeting PMIX/X could be deployed as prophylaxis to reduce the risk of resistance by limiting parasite exposure while facilitating immune education through liver stage parasite killing with each mosquito bite. This approach would facilitate CALM immunization and immune boosting of endemic populations. Humans may have a greater immune response to CALM vaccination than mice, as the liver replication phase lasts 7 or more days compared with approximately 2.5 days in mice, and therefore may allow a longer time of exposure to the immune system following boosts. Whether CALM will be efficacious as an LAI with natural exposure is worthy of further investigation. It may also be investigated as a single-shot IV vaccine similar to GAPs with the advantage of attenuating all *Plasmodium* species for which sporozoites can be readily produced, including *P. vivax*. The development of PMIX/X dual inhibitors is therefore of high importance for addressing different target product profiles that could provide protection against different forms of malaria.

## Materials and Methods

### Mice

Female and male BALB/c, C57BL/6 and ASMU-Swiss Outbred mice (4-8 weeks of age) were kept in individually ventilated cages or exhausted ventilated cages with corncob bedding, but otherwise under standard conditions with 22°C and 40-70% relative humidity, Barastock Rat & Mouse pellet (WEHI Mouse Breeder Cubes, Irradiated [#102119] and Mouse Custom Mash Irradiated [#102121], Ridley Agriproducts Pty Ltd) and water *ad libitum*. There was a maximum of 6 adults per cage. Water was filtered and acidified to pH3 and supplied in Hydropac pouch or water bottles. Animal enrichment (eg, tissues, domes) were provided weekly. Animal technicians were responsible for the daily husbandry of the mice and serviced each cage at least twice a week. Cages were cleaned on a rotating schedule (depending on the stocking density) or as often as needed. Typically, cages of 4 to 6 mice required fortnightly cleaning and cages housing 1 to 3 mice required monthly cleaning. FRG huHep mice (both male and female) were purchased from Yecuris, Inc. Animals received 3.25% dextrose type II sterile drinking water containing 8 µg/mL of NTBC on days 1 to 3 to block toxic metabolite accumulation, followed by 3.25% dextrose water on days 4 to 10 and 18 to 24, and 3.25% dextrose containing 125 µg/mL enrofloxacin antibiotics (Baytril) on days 11-17 and 25-28, with this 28-day cycle repeated throughout the animal’s lifetime. The use of animals was approved by the Walter and Eliza Hall Institute of Medical Research Animal Ethics Committee under approval number 2021.064.

### Mosquitoes

*Anopheles stephensi* mosquitoes were reared and maintained in the insectary at the Walter and Eliza Hall Institute of Medical Research according to standard methods. The mosquitoes were stored in BugDorm insect cages, mixed genders with an environment of 26°C to 27°C, relative humidity of 70% to 80%, light cycle 12 h/12 h including 30 min of ramping up or down between dark and light to imitate dawn and dusk. They were fed on reverse osmosis filtered water via cotton wicks and sugar cubes (sucrose-CSR).

### Plasmodium parasites

To produce PbmChLuc(*89*) or PbPfCSP(*59*) sporozoites for mice infection, Swiss Webster or BALB/c ‘donor’ mice were infected by intraperitoneal (IP) injection with blood stage *P. berghei* parasites. Parasitaemia was monitored by Giemsa-stained tail blood smears. Four days post IP injection, donor mice blood was transferred to naive “acceptor” mice via IP or IV injection. At ≥1% parasitaemia and exhibiting exflagellation of microgametes by microscopy at 40x magnification, acceptor mice were anesthetized with ketamine/xylazine via IP inoculation, and individually placed on top of a single container of 50 female *An. stephensi* (3–5 days old) mosquitoes, allowing feeding for 15-30 min. Midgut oocysts per mosquito were determined 14 days post-blood feeding. Salivary glands were dissected 21 days post-blood feeding from cold-anesthetized and ethanol-killed mosquitoes into Schneider’s Insect media (Sigma-Aldrich) pH 7.0 to enhance viability(*90*). Salivary glands were pelleted at 3,000*g* for 3 minutes at 4°C, crushed with plastic pestle (Scienceware 19923-0000) in a 1.5 mL tube and recentrifuged for 1 minute at 3,000 x g and crushed a second time. A 500 µL tube with 26-gauge needle hole in the bottom was lightly packed with ∼200 µL of glass wool (Supelco, silane treated 2-0411) and placed inside a standard 1.5 mL Eppendorf tube. Crushed salivary glands were passed through the glass wool filter in 150 µL aliquots and centrifuged for 120 seconds at 1200*g* and sporozoite filtrate collected in a 1.5 mL tube on ice. 10 µL of sporozoite suspension, either pure or diluted by as much as 1:20, was placed onto a haemocytometer counting chamber (Assistent, Neubauer improved) and allowed to settle for 8 min in a humidity chamber prior to counting under phase at 400x magnification (*91*).

To produce PfGFPLuci sporozoites for infection of FRG huHep mice, 7-day old female *An. stephensi* mosquitoes were fed on asynchronous gametocytes diluted to 0.6% stage V gametocytemia, via water jacketed glass membrane feeders. Mosquitoes were sugar-starved for 48 hours following feeding to enrich for blood-fed mosquitoes. Surviving mosquitoes were provided with deionized water via paper/cotton wicks and sugar cubes. Oocyst numbers were obtained from midguts dissected from cold-anesthetized and ethanol-killed mosquitoes 7 days postfeed, followed by staining with 0.1% mercurochrome. Between days 16-20 postbloodmeal, sporozoites were dissected from the mosquito salivary glands into Schneider’s Insect media.

### Infection of mice with *Plasmodium* sporozoites

Sporozoites dissected from mosquitoes were IV injected into mice in 200 μL of Schneider’s Insect Media. For infection via mosquito bites, midgut oocyst numbers were enumerated in 15-30 mosquitoes between days 14-17 postbloodmeal to determine the percentage of infected mosquitoes. Based on the percentage of mosquitoes with oocysts, 10 infected mosquitoes were selected and placed in individual feeding cups (e.g. if 83% of mosquitoes had oocysts, then 12 mosquitoes were placed in each feeding cup). Mice were anesthetized using ketamine/xylazine and placed on the feeding cups. Mosquitoes were allowed to feed on the mice for 15 minutes, with the mice being rotated between feeding cups every minute to promote mosquito probing and ensure uniform exposure to infectious bites.

### Drug preparation for oral gavage

Plasmepsin IX and X dual inhibitors WM382(*46*) and MK-7602(*65*) were dissolved in 20% DMSO/60% PG/ 20% water (v/v/v). Chloroquine (CQ) was dissolved in sterile drinking water.

### Vaccination

Radiation-attenuated sporozoites (RAS) were produced by irradiating 40,000 PbmChLuc sporozoites dissected from salivary glands with 20,000 rads using a gamma 60 Co source prior to IV injections. For CALM, mice were intravenously infected with the indicated number of PbmChLuc sporozoites followed by oral treatment with WM382 (100 mpk) or MK-7602 (500 mpk) at the indicated timepoints. For CQ CVac (CVac), mice were intravenously infected with 40,000 PbmChLuc sporozoites and treated orally with CQ 50 mpk daily starting 4 hpi, with treatment continued for 7 consecutive days. In prime-boost experiments for RAS, CALM and CVac, singly immunized mice received an intravenous homologous booster immunization with 20,000 PbmChLuc salivary gland sporozoites.

### Parasitaemia

Following sporozoite infection, parasitaemia in mice was measured from day 3 to 30 using either microscopy (Giemsa-stained thin blood smears) or an Attune NxT flow cytometer such that a drop of whole blood was collected from the tail into 200 μL of PBS. Infected blood cells were identified by plotting mCherry signal intensity against GFP signal intensity, with GFP serving as a decoy (negative control) reporter. The number of mCherry^+^/GFP^-^ events was reported per 1×10^6^ whole blood cells, with blood from uninfected mice used as a negative control. Mice were considered protected if they remained free of parasitaemia at 30 days after sporozoite infection.

### Quantification of *P. berghei* and *P. falciparum* liver and blood infection in mice by IVIS

Mice were IP injected with 200 μL XenoLight D-Luciferin (PerkinElmer, Waltham, MA) and general anaesthesia was induced by isoflurane inhalation (2% isofluorane in oxygen). Animals were imaged within 10 min of luciferin injection using the IVIS Spectrum (Xenogen, PerkinElmer) with a 21.6 cm field of view, medium binning factor and an exposure time of 3-5 min. Bioluminescence emanating from a region of interest (ROI) covering the liver region or whole animal was quantified using Living Image software version 4.7.2 (PerkinElmer) and expressed as the total flux in photons/second (p/s). Background flux was measured using an identically sized ROI placed over the lower pelvic region below the liver for liver stage infection timepoints, or over uninfected mice for blood stage infection. The background flux was expressed as the limit of detection (LOD).

### Inhibition of liver stage development assay (ILSDA)

30,000 HepG2 cells per well were infected with 5,000 freshly dissected PbmChLuc sporozoites incubated in Roswell Park Memorial Institute (RPMI) 1640 medium, 10% FBS containing immune serum from individually vaccinated mice prior to challenge (1:20 dilution) in technical triplicate, centrifuged 500*g* for 3 min and incubated at 37°C 5% CO2 for 3-4 h to allow infection. Media was replaced after 3 h, and again daily. Washed monolayers were fixed at 44-48 hpi in 4% paraformaldehyde (PFA) in 1% PBS, washed 3 times in 1x PBS and prepared in fresh 1x PBS for immunofluorecence assays.

### Immunofluorescence microscopy

Purified PbmChLuc or PbPfCSP sporozoites fixed with 4% PFA/0.0075% glutaraldehyde in 1x PBS at room temperature (RT) for 20 min and washed 3 times in 1x PBS were spotted at 25,000 sporozoites per slide. Slides were air dried, blocked in 3% BSA/1x PBS for 1 h RT. Naive and immunized mouse serum (immune serum was collected from BALB/c mice in Figure 1f, 302 days following immunization and any before challenge, diluted 1:500), anti-PbCSP 3D11 (1:1000) and anti-PfCSP 2A10 (1:1000) were prepared in 1% BSA, 1x PBS, incubated on slides for 1h RT, rinsed 3x times in 1x PBS and incubated with indicated secondary antibodies in 1% BSA/1x PBS for 1hRT before rinsing 3x times in 1x PBS, air dried and mounted with ProLong Gold Antifade (ThermoFisher). Sporozoites were imaged using the tiling function on a Zeiss LiveCell AxioObserver with 100x objective. For ILSDAs, samples were imaged on an Opera Phenix at 20x to scan entire wells of 96-well plates and the number of mCherry^+^ liver stages were quantified.

### ELISA

MaxiSorp ELISA plates (Nunc) were coated with 1mg/mL recombinant PbCSP (residues 24-319 fused to 6x His tag; Genscript) per well in carbonate buffer (16mM Na2CO3, 34 mM NaHCO3, pH9.6). After overnight incubation at 4°C, wells were washed twice in 1x PBS, 0.05% Tween-20 (PBS-T) and blocked in 5% skim milk in PBS-T for 1 h 37°C. Mouse sera were added to the ELISA plates in a 10-step 1:2 dilution series starting at 1:100. For the standard curve, anti-PbCSP monoclonal 3D11 was diluted as was mouse serum starting at 0.05 µg/mL. After 1 h incubation at 37°C, wells were washed twice in PBS-T, incubated with HRP-conjugated goat anti-mouse IgG for 1 h 37°C, washed twice in PBS-T and incubated with TMB peroxidase substrate and peroxidase substrate solution B (SeraCare) for 5 min. Finally, 1M H₃PO₄ was added to stop the reaction. OD450 was measured in a BioTek ELx405 Select CW ELISA reader. Antibody concentrations in serum were calculated by plotting the OD450 values of the serum-treated wells with OD of known standards.

### CD8^+^ T cell depletions

Following CALM vaccination, mice were IV injected with 100 μg rat anti-CD8 antibody (YTS-169) or rat isotype control (IgG2b) (WEHI antibody facility) one day before challenge with sporozoites. CD8^+^ T cell depletion was confirmed by flow cytometry analysis of the liver, spleen and draining lymph nodes of mice as previously described (*92*).

### Tissue processing and immunophenotyping

Mouse livers were collected into RPMI 1640 media, 2% FCS with 10 U heparin. Livers were passed through 100 μm strainers, washed in media and resuspended in 30 mL 35% Percoll (GE Health Care) before centrifugation 500*g* 20 min, 20°C to 22°C with no brake. Pellets were incubated in 5 mL erythrocyte lysis solution for 5 min at 20°C to 22 °C, before washing with 30 mL RPMI 1640 and resuspension in 1 mL fluorescence activated cell sorting (FACS) buffer (PBS, 5% w/v BSA, 5mM EDTA) for flow cytometry. One quarter of the cell suspension was used for antibody staining. Spleen and lymph node lymphocytes were isolated by teasing tissue through a 40 µm strainer, washing in RPMI 1640, 2% FCS, incubating cells in erythrocyte lysis solution for 1-2 min at 20-22°C, washing again and resuspending in FACS buffer. Approximately 1/25th of final volume was used for antibody staining for Fc block (1:500; BD Cat#553142), CD3 – BV711 (1:200; BD Horizon Cat#563123, clone: 145-2C11), CD4 – BV421 (1:500; BD Horizon Cat#562891, clone: GK1.5), CD8 – BV510 (1:500; BD Horizon Cat#563068, clone: 53-6.7), CD19 – PerCP (1:200; BD Pharmingen Cat#551001, clone: 1D3), CD44 (IM7; BD Biosciences), CD8α (53–6.7), CXCR6 (SA051D1), CX3CR1 (SA011F11), CD19 (1D3/CD19), CD62L (MEL-14), KLRG1 (2F1; Biolegend USA), CD69 (H1.2F3; ThermoFisher Scientific). Dead cells were excluded by propidium iodide (PI) staining (1:2000). Lymphocytes were stained with tetramers for 1 hour at 20-22 °C before staining with surface antibodies. Samples were run on an LSRFortessa (BD Biosciences) and data were analyzed using Flowjo software (TreeStar).

### *P. falciparum* liver and blood infection in FRG huHep mice

FRG huHep mice were IV infected with 300,000 *P. falciparum* NF54 sporozoites expressing GFP-luciferase(*72*) sporozoites. Mice received 200 µL 1x penicillin/streptomycin (ThermoFisher) IP on days 1, 3, 5, 6 postinfection. On day 6 postinfection, mice were injected with 50 μL clodronate liposomes IV and IP to eliminate blood and intraperitoneal phagocytic cells, respectively, and 150mpk cyclophosphamide IP to deplete neutrophils. To enable liver-to-blood transition of egressing liver merozoites, mice were IP injected with 700 µL human red blood cells (RBC) at 70% hematocrit in RPMI 1640 media on day 6 postinfection. On days 6 and 7 postinfection, mice were IV injected with 250 µL human RBC at 70% hematocrit in RPMI 1640 media. Blood was drawn from mice by retro-orbital bleeds and transferred to *in vitro* culture on day 7.7 and cultured in in RPMI 1640 medium containing 25 mM HEPES, 0.2% sodium bicarbonate and 10% human serum in the presence of fresh type O^+^ human erythrocytes at 4% hematocrit. Cultures were incubated at 37°C in a 94% N, 5% CO2, 1% O2 atmosphere for up to 8 weeks. *P. falciparum* liver and blood stage infection was measured by IVIS as above.

## Quantification and statistical analysis

All graphs of experimental data and statistical analyses were generated with GraphPad Prism 10. Statistical details of experiments are indicated in the Figure legends. \**P*<0.05, \*\**P*<0.01, \*\*\**P*<0.001. A *P* value >0.05 was considered not significant.

---

**Fig. S1 Extended analysis of BALB/c mouse protection by WM382 CALM. (A)** Liver bioluminescence signals in all mice from Fig. 1D (challenged 95-167 days post-WM382 CALM vaccination) 48 hours postchallenge (hpc). Circles represent individual mice. Dashed line is mean signal of the pelvic region of all mice and the limit of detection (LOD). Mean ± SD. Data analyzed by one-way ANOVA with Dunn multiple comparisons test; \**P*<0.05, \*\*\**P*<0.001. **(B)** Liver bioluminescence signals in all mice from Fig. 1F, G (challenged 303 to 643 days post-WM382 or CQ treatment) 48 hours postchallenge (hpc). Circles represent individual mice. Dashed line is mean signal of the pelvic region of all mice and the limit of detection (LOD). Mean ± SD. Data analyzed by one-way ANOVA with Tukey multiple comparisons test; \**P*<0.05, \*\**P*<0.01. **(C)** Protection of mice in Fig. 1D (challenged on day 167) from blood stage infection for 30 days. All mice previously treated with WM382 (5/5) had sterile protection. The five surviving mice were rechallenged by mosquito bites 367 days following vaccination and four of five mice remained protected for 30 days. The four surviving mice were rechallenged by mosquito bites 583 days following vaccination and retained sterile protection for 30 days.

**Fig. S2 WM382 CALM vaccination with low-dose sporozoite immunization. (A)** IVIS imaging of liver parasite bioluminescence 48 hours postinfection (hpi) with 500, 1,000, or 40,000 sporozoites IV with PbmChLuc sporozoites per mouse. **(B)** Mice from (A) were dosed with WM382 2×100 mg/kg (mpk) at 36 and 48 hpi) curing all mice by preventing liver-to-blood transition (n=3 mice per group). **(C)** Sporozoite infection and WM382 treatment (see B) vaccinates mice against bites from 10 PbmChLuc infectious mosquitoes, but not naive mice, 95 days later, measured 48 h postchallenge (hpc) by IVIS. **(D)** Liver bioluminescence signals in all 12 mice from (C) 48 hpc. Circles represent individual mice. Protected mice surviving after day 30 (see E) and new naive controls were rechallenged by bites of 10 PbmChLuc mosquitoes on day 173 and liver bioluminescence in all mice is shown. Protected mice surviving after day 30 (see E) and new naive controls were rechallenged as above on days 363 and 740 following vaccination and monitored for 30 days. Data for challenge 567 days following vaccination (see **e**) were not collected due to servicing of IVIS machine. Dashed line is mean signal of the pelvic region of all mice and the limit of detection (LOD). Mean ± SD. Data analyzed by one-way ANOVA with Holm-Šídák multiple comparisons test; ns, not significant; \*\**P*<0.01, \*\*\**P*<0.001. **(E)** Survival curves show percentage of mice from (D) protected against bites from 10 PbmChLuc mosquitoes at days 95, 173, 363, 567, and 740 following vaccination after 30 days per challenge. Data analyzed by log-ranked test; *P*<0.05 was significant. **(F)** Percentage of mice from (D, E) with sterile (white) or not protected (black) against bites from 10 PbmChLuc mosquitoes, measured by blood parasitemia for 30 days. Numbers above each bar show the number of protected out of the total mice challenged, derived from one independent experiment (n=3 mice per group) in which surviving mice were rechallenged (for a total of four to five challenges). Note that only two of three mice that survived the 363-day challenge (1,000 SPZ group) and one of three mice that survived the 567-day challenge (40,000 SPZ group) were subsequently rechallenged on day 567 or day 740, respectively, as the other mice were euthanized due to age-related (nonmalaria) humane endpoints.

**Fig. S3 Extended analysis of humoral and cellular responses in BALB/c mice following WM382 CALM. (A)** Technical replicate experiment for Fig. 2E, showing number of PbmChLuc liver stages per well after incubation of sporozoites with immune serum from CALM or CVac mice (1:20 dilution) or control antibodies. Circles represent serum from individually vaccinated mice (n=3-5 mice). Mean ± SD. Data analyzed by one-way ANOVA with Holm-Šídák multiple comparisons test or unpaired *t*-test; \*\*\*\**P*<0.0001. **(B-C)** Representative experiment showing the number of PbmChLuc (B) or PbPfCSP (C) liver stage parasites per well after incubation of sporozoites with pooled immune serum from previously vaccinated mice (1:20 dilution) or control antibodies. Circles represent technical replicates for each condition. Mean ± SD. Data analyzed by one-way ANOVA with Holm-Šídák multiple comparisons test or unpaired *t*-test; ns, not significant; \*\**P*<0.01, \*\*\**P*<0.001. **(D)** Quantification of CD8^+^ T cells in the liver, spleen and draining lymph nodes from BALB/c mice following IP injection of isotype control or anti-CD8 depleting antibodies 94 days after WM382 CALM a single vaccination (related to Fig. 2H-J and S3E-G). **(E)** WM382 CALM vaccinated BALB/c mice were IP treated with isotype control or anti-CD8 depleting antibodies 94 days later and challenged with 10,000 sporozoites IV 1 day later. Representative IVIS imaging shows parasite liver loads 48 hours postchallenge (hpc). **(F)** Liver bioluminescence signals in all mice from (E) 48 hpc. Circles represent individual mice. Dashed line is mean signal of the pelvic region of all mice and the limit of detection (LOD). Mean ± SD. Data analyzed by one-way ANOVA with Holm-Šídák multiple comparisons test; \*\*\**P*< 0.001. **(G)** Percentage of mice from (F) with sterile protection (white) or not protected (black) against challenge with 10,000 sporozoites IV, measured by blood parasitemia for 30 days. Numbers above bars show the number of protected out of the total mice challenged, derived from one independent experiment (n=4-5 mice per group).

**Fig. S4 WM382 attenuates *P. berghei* liver-to-blood transition in C57BL/6 mice. (A)** Mice were infected with 40,000 PbmChLuc sporozoites IV and given WM382 orally 36- and 48-hours postinfection (hpi). From 52-70 hpi, bioluminescence was measured to follow liver infection and egress, while flow cytometry measured the first wave of blood infection. **(B)** Bioluminescent images show peak liver infection (52 hpi), liver egress (55 hpi), and blood infection (70 hpi) in WM382-treated and untreated mice (n=12 mice per group). **(C)** Liver infection (52 hpi) was similar between treatment groups and significantly reduced at 55 hpi. Circles represent individual mice. Mean ± SD; \*\*\**P*<0.001. **(D)** Percentage loss of bioluminescence between 52 and 55 hpi (representative of liver egress). Mean ± SD; \**P*<0.05. **(E)** Mouse whole-body luminescence at 70 hpi (indicative of blood infection). Mean ± SD; \*\*\**P*<0.001. **(F)** Blood parasitemia (iRBCs per million cells) in mice determined by flow cytometry of mCherry signals at 70 hpi. Mean ± SD. \*\*\**P*<0.001. **(G)** Time to blood stage infection in mice from (E, F). Data are pooled from n=2 independent experiments. In (c, d), dashed line is mean signal of the pelvic region of all mice and the limit of detection (LOD).

**Fig. S5 WM382 CALM prime and boost vaccinations of B6 mice and liver Trm analyses. (A)** B6 mice given a WM382 CALM primary vaccination (CALM P) were challenged months later by bites of 10 infectious mosquitoes. **(B)** CALM P generates nonsterilizing pre-erythrocytic immunity in B6 mice when challenged 108 to 133 days postvaccination as in (A), measured 48 hours postchallenge (hpc) by IVIS. **(C)** Liver bioluminescence signals in all mice from (B) at 48 hours postchallenge (hpc). Circles represent individual mice. Dashed line is mean signal of the pelvic region of all mice and the limit of detection (LOD). Mean ± SD; n=7-16 mice per group; data analyzed by unpaired *t*-test; \*\**P*<0.01. **(D)** Time to patent blood infection for all mice shown in (c). Data are pooled from n=2 independent experiments. **(E)** Numbers of memory CD8^+^ T cells in livers of B6 mice immunized with radiation-attenuated sporozoites (RAS), WM382 CALM or CQ CVac prime (P) and prime-boost regimens (PB; same data as in Fig. 5G) quantified by flow cytometry 42 days after the boost immunization. Naive (Tnaive; CD44^-^ CD8^+^), tissue resident memory (Trm; CD44^+^ CD69^+^ CD62L^-^), effector memory (Tem; CD44^+^ CD69^-^ CD62L^-^), and central memory (Tcm; CD44^+^ CD69^-^ CD62L^+^) T cells shown. Counts are mean ± SEM compared by one-way ANOVA with Tukey’s multiple comparisons test; \*\**P*<0.01, \*\*\**P*<0.001, \*\*\*\**P*<0.0001. **(F)** Representative FACS plots of memory CD8^+^ T cells isolated from mice (Fig. 5H-K) specific to *P. berghei* SERA1, RPL6, RPA1, and TRAP in the liver.

**Fig. S6 Chemoprotection and CALM vaccination with WM382 in Swiss outbred mice. (A)** Infection of Swiss outbred mice with 40,000 PbmChLuc sporozoites and treatment with WM382 orally at 36- and 48-hours postinfection or chloroquine (CQ) orally for 7 days, prevents blood stage infection, for 30 days (chemoprotection). This event is considered a CALM prime (CALM P) or CQ CVac prime (CQ CVac P) immunization, respectively. **(B)** Mice from (B) that received CALM P (n=4 mice) or CQ CVac P (n=5 mice) were challenged 12 weeks later with the bites of 10 PbmChLuc mosquitoes and protection from blood stage infection was followed for 30 days. **(C)** Mice from (A) that received CALM (n=5 mice) or CQ CVac (n=5 mice) were given a boost vaccination (prime-boost; PB) 6 weeks later and challenged a further 6 weeks later with the bites of 10 PbmChLuc infectious mosquitoes and protection from blood stage infection was followed for 30 days. **(D)** Liver bioluminescence signals in all mice from (A-C), challenged 12 weeks (83 days) following prime vaccination, measured 49 hours postchallenge (hpc). Circles represent individual mice. Dashed line is mean signal of the pelvic region of all mice and the limit of detection (LOD). Mean ± SD. Data analyzed by one-way ANOVA with Holm-Šídák multiple comparisons test; \*\*\**P*<0.001. **(E-F)** Time to patent blood infection for all mice shown in (A-D) given WM382 CALM prime (P) and prime-boost (PB) (E) or CQ CVac prime (P) and prime-boost (PB) (F) vaccinations.

**Fig. S7 WM382 CALM primary vaccination stimulates moderate functional CD8^+^ T cell responses in B6 mice. (A)** B6 mice given a primary WM382 CALM vaccination (CALM P) were, 15 weeks (107 days) later, injected IP with isotype control or anti-CD8 depleting antibodies and challenged by IV injection with 10,000 PbmChLuc sporozoites 1 day later. **(B)** FACS quantification of CD8^+^ T cells in the liver of mice from (A) or naive controls one day following IP injection of isotype control or anti-CD8 depleting antibodies. **c,** IVIS imaging of mice from (A) shows parasite liver loads 48 hours postchallenge (hpc). **(D)** Liver bioluminescence signals in all mice from panel (c) 48 hours postchallenge. Circles represent individual mice. Dashed line is mean signal of the pelvic region of all mice and the limit of detection (LOD). Mean ± SD. Data from one independent experiment (four mice per group) were analyzed by one-way ANOVA with Dunnett multiple comparisons test; ns, not significant; \**P*<0.05. **(E)** Time to blood-stage infection for all CALM mice in (D).

**Fig. S8 Depletion of CD8^+^ T cells following WM382 CALM prime-boost vaccination of B6 mice.** FACS quantification of CD8^+^ T cells in the liver, spleen, and draining lymph nodes of mice from Fig. 5C-F 1 day following IP injection of isotype control or anti-CD8-depleting antibodies.

## ACKNOWLEDGEMENTS

We thank the Melbourne Red Cross for human erythrocytes, Waail Abdalla, Cody Allison, Danielle Boyd, Gregor Ebert, and Robyn McConville for technical support with mouse studies, Volker Huessler for *P. berghei* expressing mCherry and luciferase, Fidel Zavala for *P. berghei* expressing PfCSP and for PfCSP antibodies, and Stefen Kappe for *P. falciparum* NF54 expressing GFP-luciferase. Hybridoma 3D11 anti-*P. berghei* 44-Kilodalton Sporozoite Surface Protein (Pb44) was obtained from BEI Resources (MRA-100), contributed by Victor Nussenzweig. *Anopheles stephensi* mosquitoes were originally imported into the WEHI Insectary from Johns Hopkins School of Public Health, USA. All data in this study are free to access and use. **Funding:** This work was possible through funding from The Wellcome Trust (202749/Z/16/Z, 219658/Z/19/Z), MSD Sponsored Research Agreement (LKR219658), the National Health and Medical Research Council of Australia (NHMRC; JAB is a Leadership Fellow, 1176955) and supported by the Victorian State Government Operational Infrastructure Support grant (Institutional grant) and Australian Government NHMRC IRIISS. **Author Contributions:** R.W.J.S., W.R.H., D.B.O., and J.A.B. conceptualized the studies. J.A.B., R.W.J.S., and Y.C.C. wrote the draft, and all authors contributed to data analysis, data interpretation, reviewing and editing the manuscript. R.W.J.S., Y.C.C., S.C., and D.F-R. performed biological, mouse, and immunological experiments. E.H. and J.A.B. performed transcriptomic and proteomic analyses. J.A.B. supervised the study.

## Data and materials availability

Further information and requests for resources and reagents should be directed to and will be fulfilled by the Lead Contact, Justin A. Boddey. All unique/stable reagents generated in this study are available from the Lead Contact with a completed Materials Transfer Agreement.

## REFERENCES AND NOTES

1. WHO, “Malaria Facts Sheet,” (2024).

2. A. M. Vaughan, S. H. Kappe, Malaria Parasite Liver Infection and Exoerythrocytic Biology. Cold Spring Harb Perspect Med, (2017).

3. A. Sturm et al., Manipulation of host hepatocytes by the malaria parasite for delivery into liver sinusoids. Science 313, 1287–1290 (2006).

4. K. Baer, C. Klotz, S. H. Kappe, T. Schnieder, U. Frevert, Release of hepatic Plasmodium yoelii merozoites into the pulmonary microvasculature. PLoS Pathog 3, e171 (2007).

5. S. C. T. P. RTS, Efficacy and safety of RTS,S/AS01 malaria vaccine with or without a booster dose in infants and children in Africa: final results of a phase 3, individually randomised, controlled trial. Lancet 386, 31–45 (2015).

6. A. Dicko et al., Seasonal vaccination with RTS,S/AS01(E) vaccine with or without seasonal malaria chemoprevention in children up to the age of 5 years in Burkina Faso and Mali: a double-blind, randomised, controlled, phase 3 trial. Lancet Infect Dis 24, 75–86 (2024).

7. M. S. Datoo et al., Safety and efficacy of malaria vaccine candidate R21/Matrix-M in African children: a multicentre, double-blind, randomised, phase 3 trial. Lancet 403, 533–544 (2024).

8. M. S. Datoo et al., Efficacy of a low-dose candidate malaria vaccine, R21 in adjuvant Matrix-M, with seasonal administration to children in Burkina Faso: a randomised controlled trial. Lancet 397, 1809–1818 (s2021).

9. D. Chandramohan et al., Seasonal Malaria Vaccination with or without Seasonal Malaria Chemoprevention. N Engl J Med 385, 1005–1017 (2021).

10. WHO, Malaria vaccines: preferred product characteristics and clinical development considerations. Geneva. Licence: CC BY-NC-SA 3.0 IGO, (2022).

11. R. A. Seder et al., Protection against malaria by intravenous immunization with a nonreplicating sporozoite vaccine. Science 341, 1359–1365 (2013).

12. M. Roestenberg et al., A double-blind, placebo-controlled phase 1/2a trial of the genetically attenuated malaria vaccine PfSPZ-GA1. Sci Transl Med 12, (2020).

13. B. C. van Schaijk et al., A genetically attenuated malaria vaccine candidate based on P. falciparum b9/slarp gene-deficient sporozoites. eLife 3, (2014).

14. S. A. Mikolajczak et al., A next-generation genetically attenuated Plasmodium falciparum parasite created by triple gene deletion. Mol Ther 22, 1707–1715 (2014).

15. J. G. Kublin et al., Complete attenuation of genetically engineered Plasmodium falciparum sporozoites in human subjects. Sci Transl Med 9, (2017).

16. Y. S. Goh, D. McGuire, L. Renia, Vaccination With Sporozoites: Models and Correlates of Protection. Front Immunol 10, 1227 (2019).

17. T. L. Richie et al., Sporozoite immunization: innovative translational science to support the fight against malaria. Expert Rev Vaccines 22, 964–1007 (2023).

18. P. E. Duffy, J. P. Gorres, S. A. Healy, M. Fried, Malaria vaccines: a new era of prevention and control. Nature reviews. Microbiology 22, 756–772 (2024).

19. D. Goswami, N. K. Minkah, S. H. I. Kappe, Designer Parasites: Genetically Engineered Plasmodium as Vaccines To Prevent Malaria Infection. J Immunol 202, 20–28 (2019).

20. S. Borrmann, K. Matuschewski, Protective immunity against malaria by ‘natural immunization’: a question of dose, parasite diversity, or both? Curr Opin Immunol 23, 500–508 (2011).

21. R. J. Nevagi, M. F. Good, D. I. Stanisic, Plasmodium infection and drug cure for malaria vaccine development. Expert Rev Vaccines 20, 163–183 (2021).

22. N. S. Butler et al., Superior antimalarial immunity after vaccination with late liver stage-arresting genetically attenuated parasites. Cell Host Microbe 9, 451–462 (2011).

23. D. Goswami et al., A replication competent Plasmodium falciparum parasite completely attenuated by dual gene deletion. EMBO Mol Med 16, 723–754 (2024).

24. B. Franke-Fayard et al., Creation and preclinical evaluation of genetically attenuated malaria parasites arresting growth late in the liver. NPJ Vaccines 7, 139 (2022).

25. O. A. C. Lamers et al., Safety and Efficacy of Immunization with a Late-Liver-Stage Attenuated Malaria Parasite. N Engl J Med 391, 1913–1923 (2024).

26. G. V. T. Roozen et al., Single immunization with genetically attenuated PfΔmei2 (GA2) parasites by mosquito bite in controlled human malaria infection: a placebo-controlled randomized trial. Nat Med 31, 218–222 (2025).

27. A. Mishra, P. Paul, M. Srivastava, S. Mishra, A Plasmodium late liver stage arresting GAP provides superior protection in mice. NPJ Vaccines 9, 193 (2024).

28. J. E. Epstein et al., Live attenuated malaria vaccine designed to protect through hepatic CD8(+) T cell immunity. Science 334, 475–480 (2011).

29. I. A. Cockburn et al., In vivo imaging of CD8+ T cell-mediated elimination of malaria liver stages. Proc Natl Acad Sci U S A 110, 9090–9095 (2013).

30. L. Schofield et al., Gamma interferon, CD8+ T cells and antibodies required for immunity to malaria sporozoites. Nature 330, 664–666 (1987).

31. F. N. Watson et al., Cryopreserved Sporozoites with and without the Glycolipid Adjuvant 7DW8-5 Protect in Prime-and-Trap Malaria Vaccination. The American journal of tropical medicine and hygiene 106, 1227–1236 (2022).

32. D. Fernandez-Ruiz et al., Liver-Resident Memory CD8(+) T Cells Form a Front-Line Defense against Malaria Liver-Stage Infection. Immunity 45, 889–902 (2016).

33. A. S. Ishizuka et al., Protection against malaria at 1 year and immune correlates following PfSPZ vaccination. Nat Med 22, 614–623 (2016).

34. S. W. Tse, A. J. Radtke, D. A. Espinosa, I. A. Cockburn, F. Zavala, The chemokine receptor CXCR6 is required for the maintenance of liver memory CD8(+) T cells specific for infectious pathogens. J Infect Dis 210, 1508–1516 (2014).

35. A. Noe et al., Deep Immune Phenotyping and Single-Cell Transcriptomics Allow Identification of Circulating TRM-Like Cells Which Correlate With Liver-Stage Immunity and Vaccine-Induced Protection From Malaria. Front Immunol 13, 795463 (2022).

36. E. Belnoue et al., Protective T cell immunity against malaria liver stage after vaccination with live sporozoites under chloroquine treatment. J Immunol 172, 2487–2495 (2004).

37. E. M. Bijker et al., Cytotoxic markers associate with protection against malaria in human volunteers immunized with Plasmodium falciparum sporozoites. J Infect Dis 210, 1605–1615 (2014).

38. M. N. de Menezes et al., Long lived liver-resident memory T cells of biased specificities for abundant sporozoite antigens drive malaria protection by radiation-attenuated sporozoite vaccination. PLoS Pathog 21, e1012731 (2025).

39. B. Mordmuller et al., Sterile protection against human malaria by chemoattenuated PfSPZ vaccine. Nature 542, 445–449 (2017).

40. Z. Sulyok et al., Heterologous protection against malaria by a simple chemoattenuated PfSPZ vaccine regimen in a randomized trial. Nature communications 12, 2518 (2021).

41. A. Mwakingwe-Omari et al., Two chemoattenuated PfSPZ malaria vaccines induce sterile hepatic immunity. Nature 595, 289–294 (2021).

42. T. Sahu et al., Chloroquine neither eliminates liver stage parasites nor delays their development in a murine Chemoprophylaxis Vaccination model. Front Microbiol 6, 283 (2015).

43. J. Friesen et al., Natural immunization against malaria: causal prophylaxis with antibiotics. Sci Transl Med 2, 40ra49 (2010).

44. D. Chandramohan et al., Effect of Adding Azithromycin to Seasonal Malaria Chemoprevention. N Engl J Med 380, 2197–2206 (2019).

45. N. K. Minkah, S. H. I. Kappe, Malaria vaccine gets a parasite boost in the liver. Nature 595, 173–174 (2021).

46. P. Favuzza et al., Dual Plasmepsin-Targeting Antimalarial Agents Disrupt Multiple Stages of the Malaria Parasite Life Cycle. Cell Host Microbe 27, 642–658 e612 (2020).

47. M. de Lera Ruiz et al., The Invention of WM382, a Highly Potent PMIX/X Dual Inhibitor toward the Treatment of Malaria. ACS Med Chem Lett 13, 1745–1754 (2022).

48. M. J. Shears et al., Proteomic Analysis of Plasmodium Merosomes: The Link between Liver and Blood Stages in Malaria. J Proteome Res 18, 3404–3418 (2019).

49. R. Caldelari et al., Transcriptome analysis of Plasmodium berghei during exo-erythrocytic development. Malaria journal 18, 330 (2019).

50. P. Pino et al., A multi-stage antimalarial targets the plasmepsins IX and X essential for invasion and egress. Science, (2017).

51. A. S. Nasamu et al., Plasmepsins IX and X are essential and druggable mediators of malaria parasite egress and invasion. Science, (2017).

52. A. N. Hodder et al., Basis for drug selectivity of plasmepsin IX and X inhibition in Plasmodium falciparum and vivax. Structure 30, 947–961 e946 (2022).

53. L. Tawk et al., A key role for Plasmodium subtilisin-like SUB1 protease in egress of malaria parasites from host hepatocytes. J Biol Chem 288, 33336–33346 (2013).

54. M. Roestenberg et al., Protection against a malaria challenge by sporozoite inoculation. N Engl J Med 361, 468–477 (2009).

55. E. Belnoue et al., Regression of established liver tumor induced by monoepitopic peptide-based immunotherapy. J Immunol 173, 4882–4888 (2004).

56. J. P. Vanderberg, U. Frevert, Intravital microscopy demonstrating antibody-mediated immobilisation of *Plasmodium berghei* sporozoites injected into skin by mosquitoes. Int J Parasitol 34, 991–996 (2004).

57. Y. Flores-Garcia et al., Antibody-Mediated Protection against Plasmodium Sporozoites Begins at the Dermal Inoculation Site. mBio 9, (2018).

58. E. Aliprandini et al., Cytotoxic anti-circumsporozoite antibodies target malaria sporozoites in the host skin. Nat Microbiol 3, 1224–1233 (2018).

59. D. A. Espinosa et al., Robust antibody and CD8(+) T-cell responses induced by P. falciparum CSP adsorbed to cationic liposomal adjuvant CAF09 confer sterilizing immunity against experimental rodent malaria infection. NPJ Vaccines 2, (2017).

60. N. Yoshida, R. S. Nussenzweig, P. Potocnjak, V. Nussenzweig, M. Aikawa, Hybridoma produces protective antibodies directed against the sporozoite stage of malaria parasite. Science 207, 71–73 (1980).

61. J. Nicholas et al., Comparative analyses of functional antibody-mediated inhibition with anti-circumsporozoite monoclonal antibodies against transgenic Plasmodium berghei. Malaria journal 22, 335 (2023).

62. P. Romero et al., Cloned cytotoxic T cells recognize an epitope in the circumsporozoite protein and protect against malaria. Nature 341, 323–325 (1989).

63. N. W. Schmidt, N. S. Butler, V. P. Badovinac, J. T. Harty, Extreme CD8 T cell requirements for anti-malarial liver-stage immunity following immunization with radiation attenuated sporozoites. PLoS Pathog 6, e1000998 (2010).

64. M. R. van Dijk et al., Genetically attenuated, P36p-deficient malarial sporozoites induce protective immunity and apoptosis of infected liver cells. Proc Natl Acad Sci U S A 102, 12194–12199 (2005).

65. P. Favuzza et al., MK-7602: a novel potent multi-stage dual-targeting antimalarial. Under review, (2025).

66. U. Frevert et al., Intravital observation of *Plasmodium berghei* sporozoite infection of the liver. PLoS Biol. 3, e192 (2005).

67. A. M. Valencia-Hernandez et al., A Natural Peptide Antigen within the Plasmodium Ribosomal Protein RPL6 Confers Liver T(RM) Cell-Mediated Immunity against Malaria in Mice. Cell Host Microbe 27, 950–962 e957 (2020).

68. L. S. Lau et al., Blood-stage Plasmodium berghei infection generates a potent, specific CD8+ T-cell response despite residence largely in cells lacking MHC I processing machinery. J Infect Dis 204, 1989–1996 (2011).

69. J. C. Hafalla et al., Identification of targets of CD8(+) T cell responses to malaria liver stages by genome-wide epitope profiling. PLoS Pathog 9, e1003303 (2013).

70. Y. C. Chua et al., Temporal expression of liver-stage malaria antigens shapes vaccine efficacy. Under revision, (2025).

71. A. M. Vaughan et al., Complete *Plasmodium falciparum* liver-stage development in liver-chimeric mice. J Clin Invest 122, 3618–3628 (2012).

72. A. M. Vaughan et al., A transgenic Plasmodium falciparum NF54 strain that expresses GFP-luciferase throughout the parasite life cycle. Mol Biochem Parasitol 186, 143–147 (2012).

73. M. Roestenberg et al., Long-term protection against malaria after experimental sporozoite inoculation: an open-label follow-up study. Lancet 377, 1770–1776 (2011).

74. S. C. Murphy et al., PfSPZ-CVac efficacy against malaria increases from 0% to 75% when administered in the absence of erythrocyte stage parasitemia: A randomized, placebo-controlled trial with controlled human malaria infection. PLoS Pathog 17, e1009594 (2021).

75. Y. Antonova-Koch et al., Open-source discovery of chemical leads for next-generation chemoprotective antimalarials. Science 362, (2018).

76. S. Mukherjee, A. S. Nasamu, K. C. Rubiano, D. E. Goldberg, Activation of the Plasmodium Egress Effector Subtilisin-Like Protease 1 Is Mediated by Plasmepsin X Destruction of the Prodomain. mBio 14, e0067323 (2023).

77. T. Triglia et al., Plasmepsin X activates the PCRCR complex of Plasmodium falciparum by processing PfRh5 for erythrocyte invasion. Nature communications 14, 2219 (2023).

78. C. Suarez, K. Volkmann, A. R. Gomes, O. Billker, M. J. Blackman, The malarial serine protease SUB1 plays an essential role in parasite liver stage development. PLoS Pathog 9, e1003811 (2013).

79. E. D. Putrianti, A. Schmidt-Christensen, V. Heussler, K. Matuschewski, A. Ingmundson, A Plasmodium cysteine protease required for efficient transition from the liver infection stage. PLoS Pathog 16, e1008891 (2020).

80. R. van Schuijlenburg et al., Early Activation of Lung CD8(+) T Cells After Immunization with Live Plasmodium berghei Malaria Sporozoites. Pathog Immun 10, 46–68 (2025).

81. S. E. Lindner et al., Transcriptomics and proteomics reveal two waves of translational repression during the maturation of malaria parasite sporozoites. Nature communications 10, 4964 (2019).

82. S. E. Lindner et al., Total and putative surface proteomics of malaria parasite salivary gland sporozoites. Mol Cell Proteomics 12, 1127–1143 (2013).

83. N. S. Butler, A. M. Vaughan, J. T. Harty, S. H. Kappe, Whole parasite vaccination approaches for prevention of malaria infection. Trends Immunol 33, 247–254 (2012).

84. E. M. Bijker et al., Protection against malaria after immunization by chloroquine prophylaxis and sporozoites is mediated by preerythrocytic immunity. Proc Natl Acad Sci U S A 110, 7862–7867 (2013).

85. C. Andolina et al., Quantification of sporozoite expelling by Anopheles mosquitoes infected with laboratory and naturally circulating P. falciparum gametocytes. eLife 12, (2024).

86. S. Kanatani, D. Stiffler, T. Bousema, G. Yenokyan, P. Sinnis, Revisiting the Plasmodium sporozoite inoculum and elucidating the efficiency with which malaria parasites progress through the mosquito. Nature communications 15, 748 (2024).

87. F. Guglielmo et al., Quantifying individual variability in exposure risk to mosquito bites in the Cascades region, Burkina Faso. Malaria journal 20, 44 (2021).

88. L. G. Bekker et al., Twice-Yearly Lenacapavir or Daily F/TAF for HIV Prevention in Cisgender Women. N Engl J Med 391, 1179–1192 (2024).

89. M. Prado et al., Long-term live imaging reveals cytosolic immune responses of host hepatocytes against Plasmodium infection and parasite escape mechanisms. Autophagy 11, 1561–1579 (2015).

90. A. Roth et al., A comprehensive model for assessment of liver stage therapies targeting Plasmodium vivax and Plasmodium falciparum. Nature communications 9, 1837 (2018).

91. J. S. Armistead et al., Plasmodium falciparum subtilisin-like ookinete protein SOPT plays an important and conserved role during ookinete infection of the Anopheles stephensi midgut. Mol Microbiol 109, 458–473 (2018).

92. G. Ebert et al., Cellular inhibitor of apoptosis proteins prevent clearance of hepatitis B virus. Proc Natl Acad Sci U S A 112, 5797–5802 (2015).

