## Supplementary figures and images for "Chemovaccination with a novel antimalarial targeting the late liver stage induces durable immunity against malaria"

### Supplementary Figures

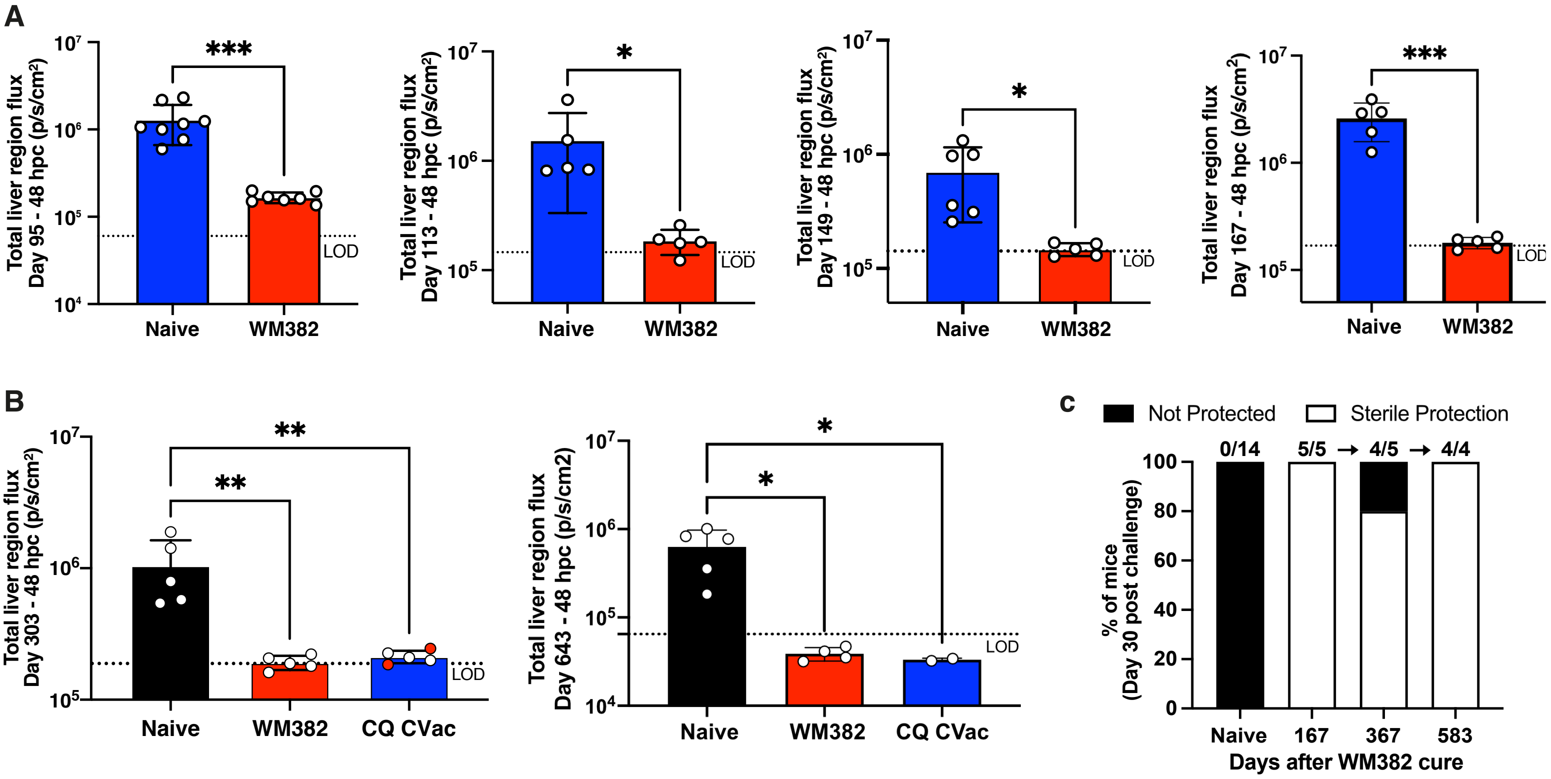

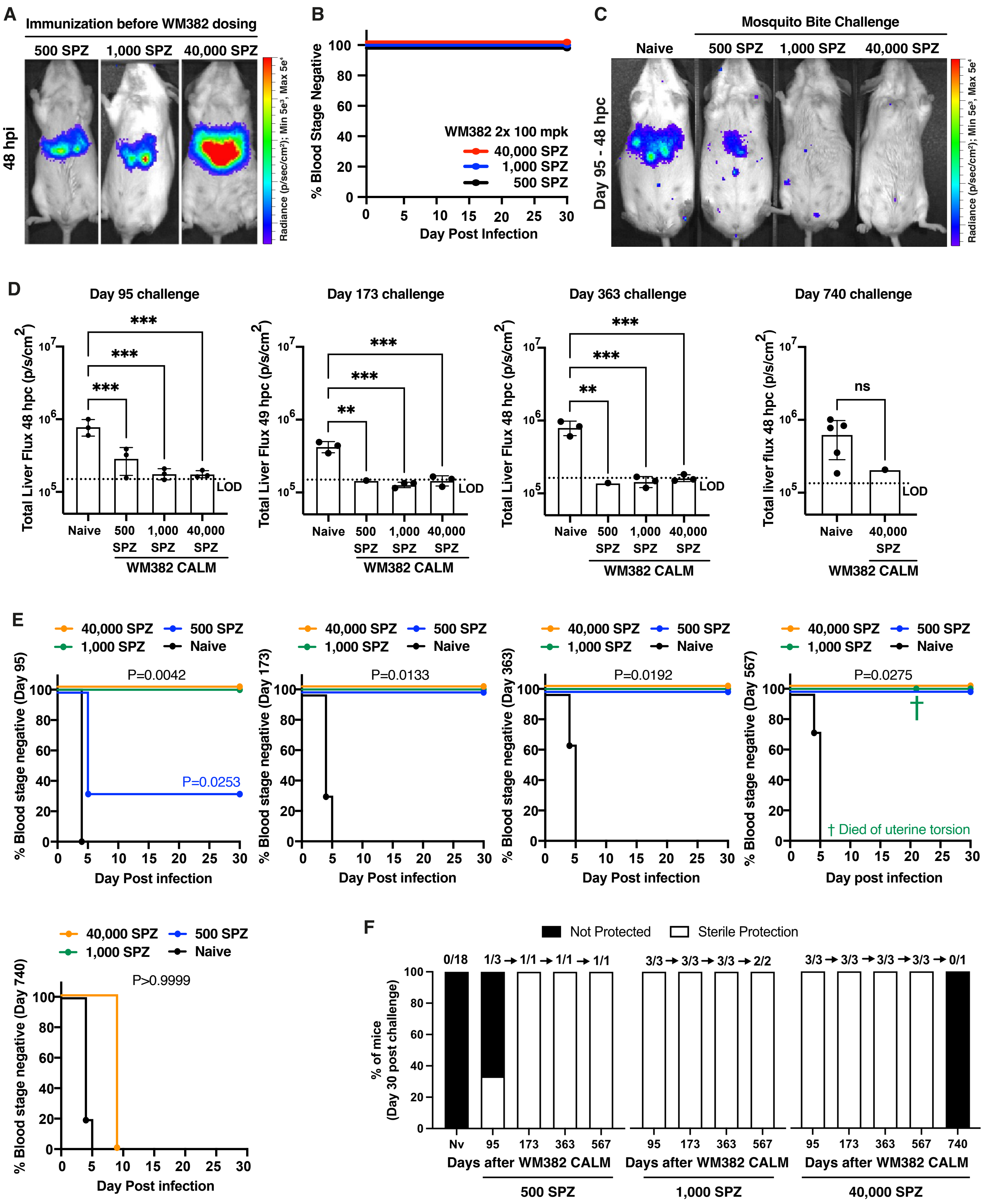

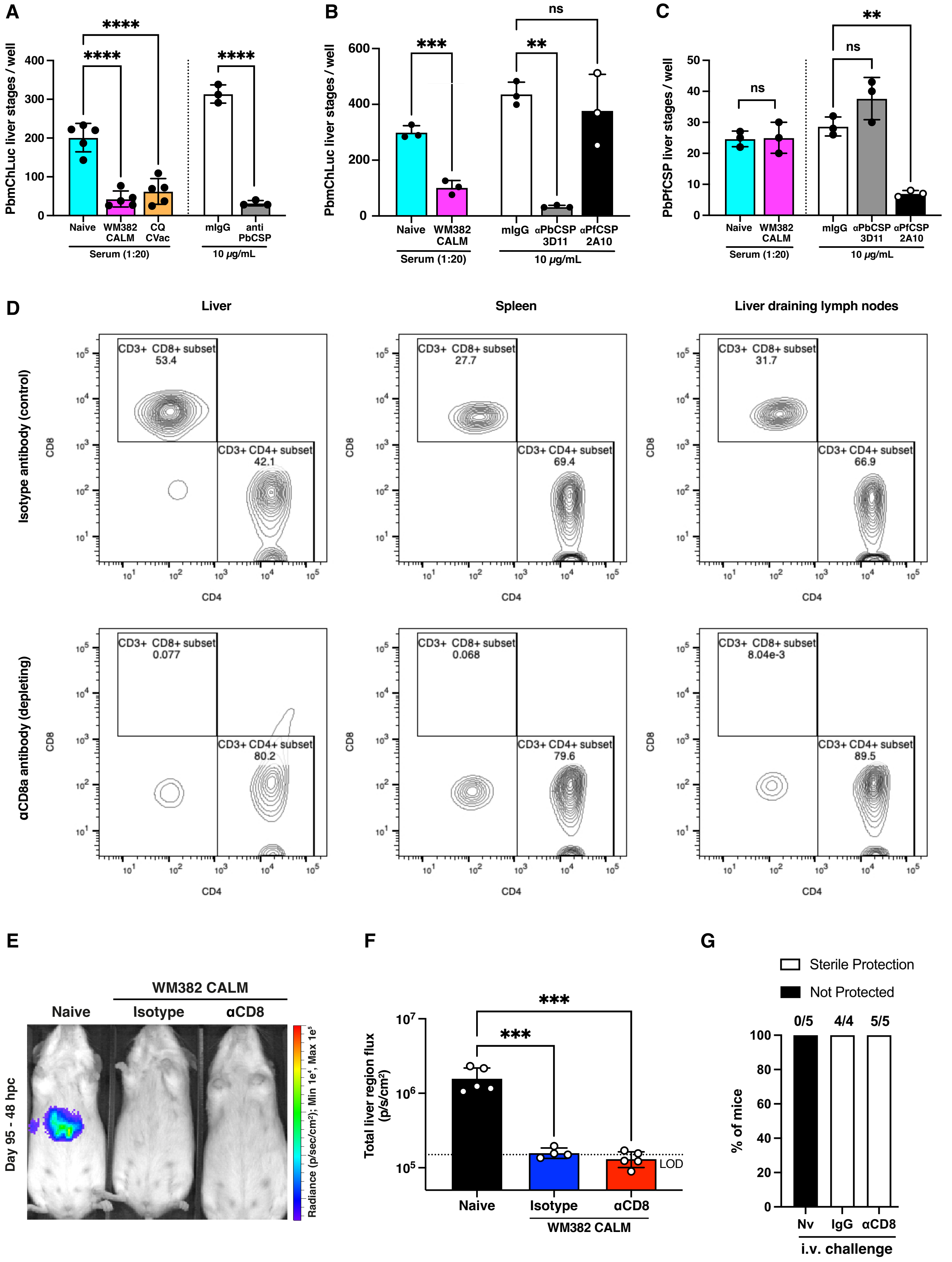

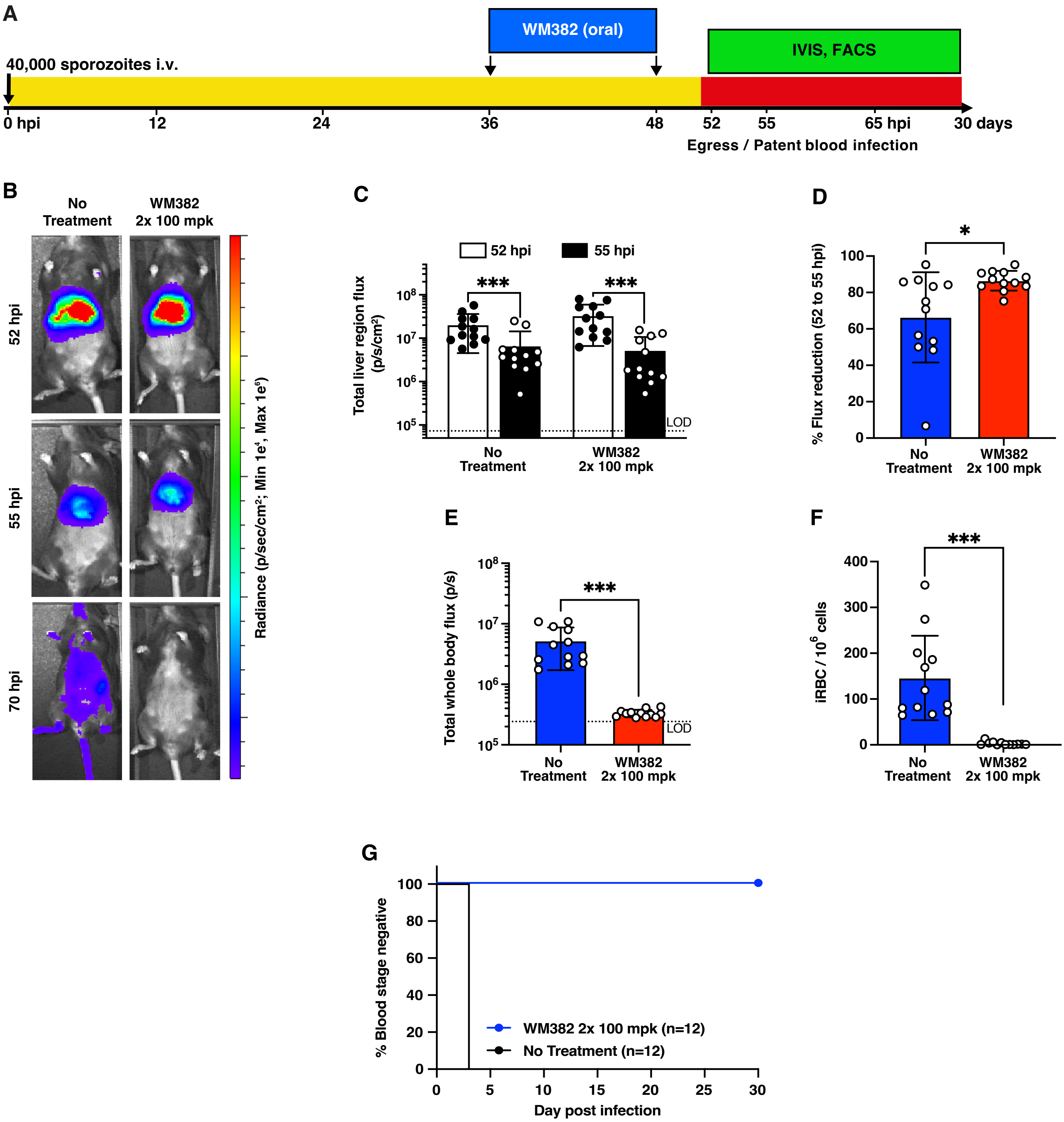

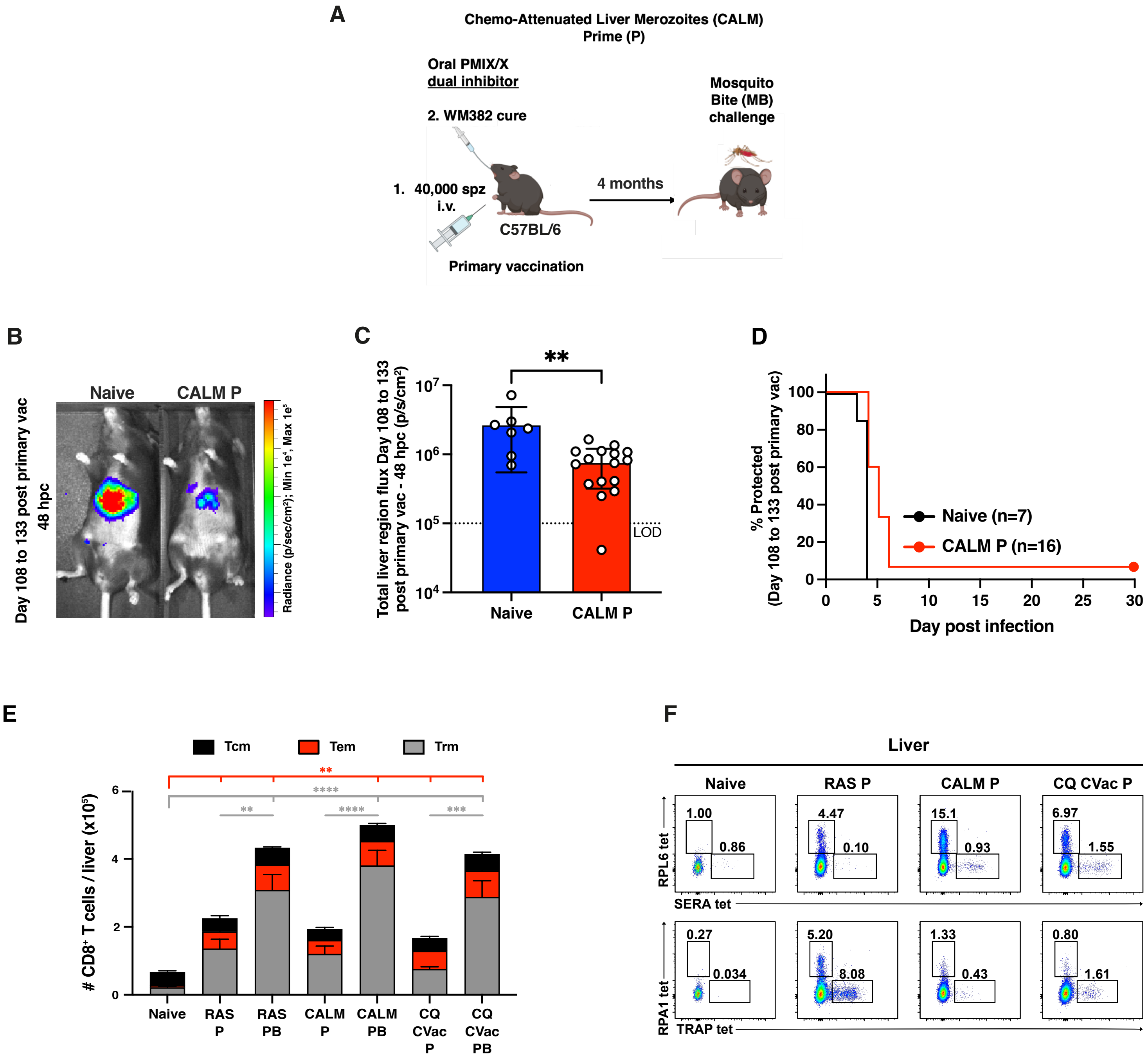

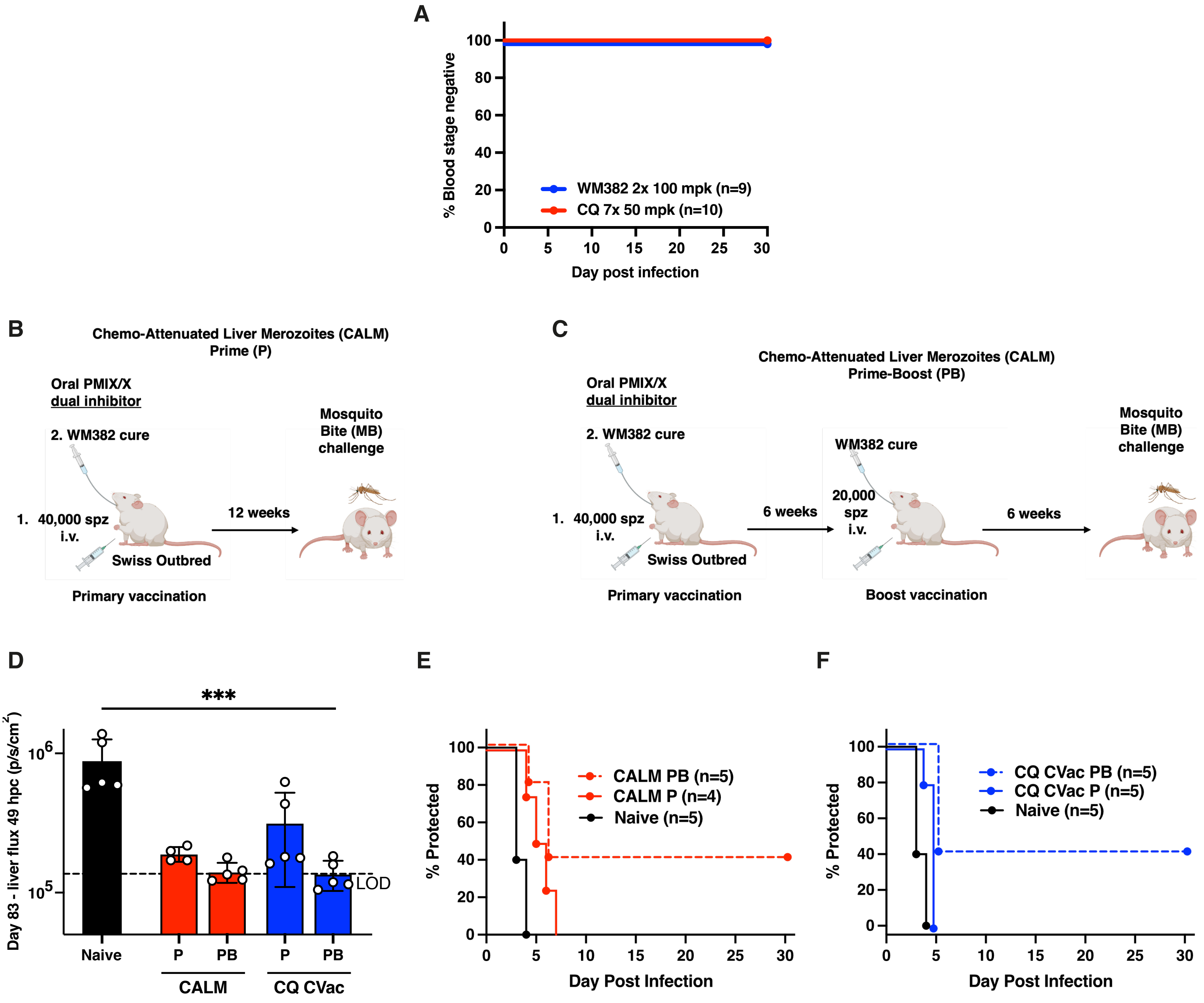

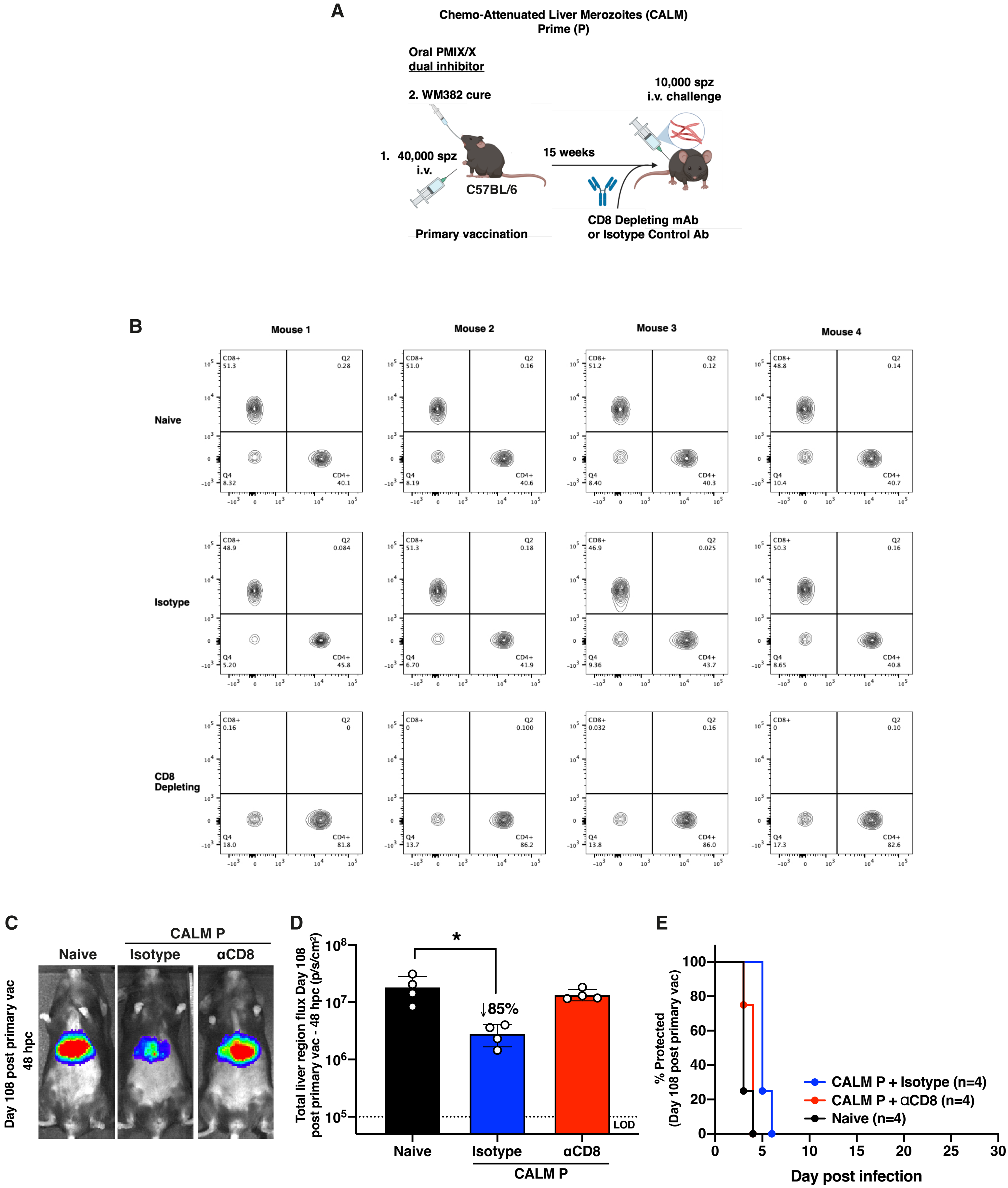

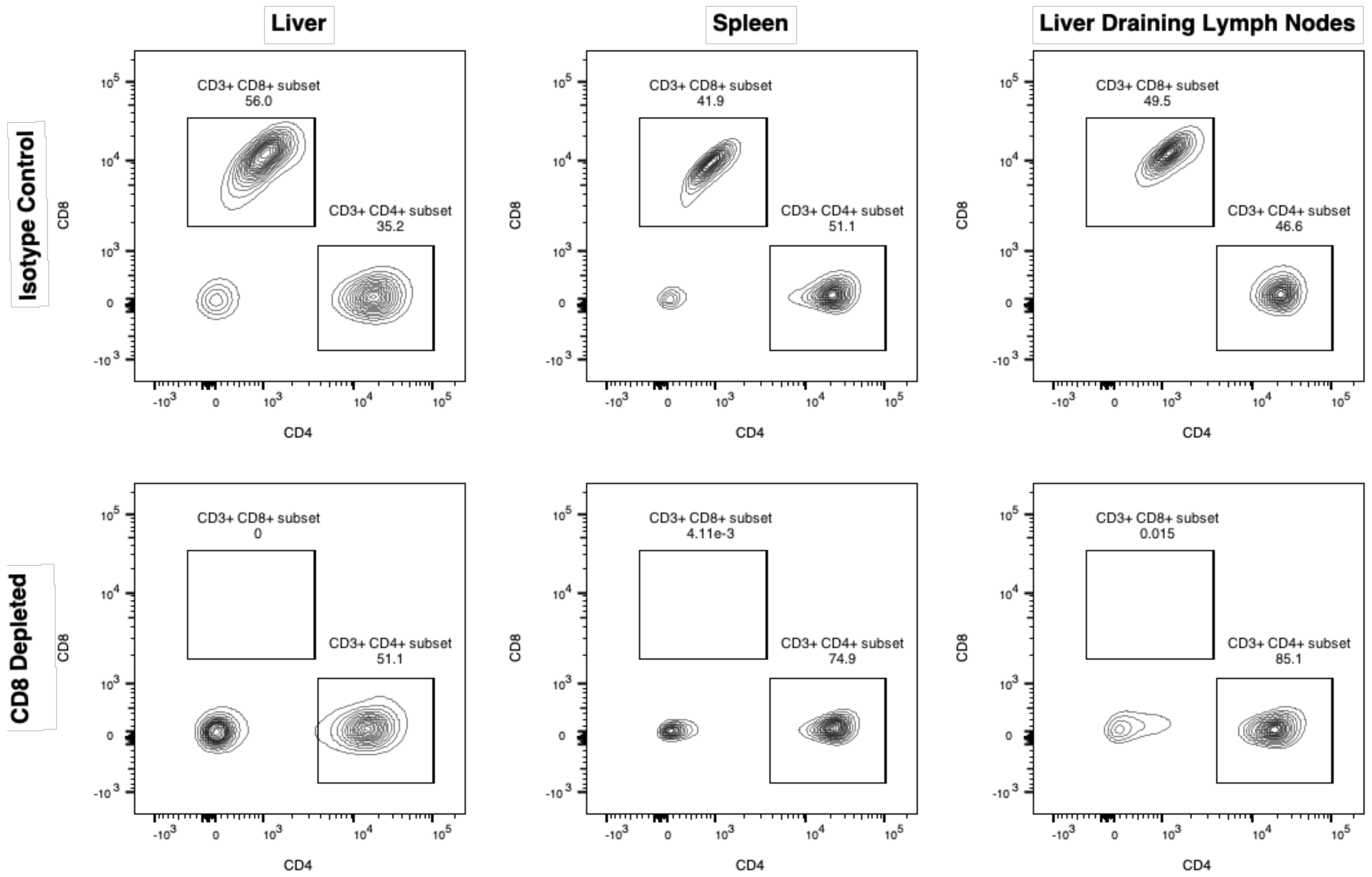
